# ESCRT-III-dependent adhesive and mechanical changes are triggered by a mechanism sensing paracellular diffusion barrier alteration in *Drosophila* epithelial cells

**DOI:** 10.1101/2023.05.24.542059

**Authors:** Thomas Esmangart de Bournonville, Mariusz K. Jaglarz, Emeline Durel, Roland Le Borgne

## Abstract

Barrier functions of proliferative epithelia are constantly challenged by mechanical and chemical constraints. How epithelia respond to and cope with disturbances of the paracellular diffusion barrier to allow tissue integrity maintenance has been poorly characterized. Cellular junctions play an important role in this process and intracellular traffic contribute to their homeostasis. Here, we reveal that, in *Drosophila* pupal notum, alteration of the bi- or tricellular septate junctions (SJs) triggers a mechanism with two prominent outcomes. On one hand, there is an increase in the levels of E-cadherin, F- Actin and non-muscle myosin II in the plane of adherens junctions. On the other hand, β-integrin/Vinculin-positive cell contacts are reinforced along the lateral and basal membranes. We report that the weakening of SJ integrity, caused by the depletion of bi- or tricellular SJ components, reduces ESCRT-III/Vps32/Shrub-dependent degradation and promotes instead Retromer-dependent recycling of SJ components. The consequence of the reduction in Shrub-dependent degradation extends to other transmembrane protein cargoes. Consequently, this trigger increased levels of β- integrin, Crumbs and the Crumbs effectors β-Heavy Spectrin Karst. We propose a mechanism by which epithelial cells, upon sensing alterations in the paracellular diffusion barrier, target Shrub to adjust the degradation/recycling balance and thereby compensate for barrier defects while maintaining epithelial integrity.

## Introduction

Epithelia are key tissues of organisms, facing the outside and protecting the inner part of the organism against both physical and chemical injuries. Although the epithelial cells composing the tissue need to establish solid and resistant barriers, they remain highly plastic. Indeed, throughout development, epithelial cells undergo profound changes in cell shape or cell–cell contacts during cell movements, divisions, cell intercalation or extrusion [1–4]. Most of these mechanisms imply junctional remodelling and rely on a set of molecular actors that form the cellular cytoskeleton, such as non-muscle myosin II (Myo-II) and filamentous actin (F-actin). Both of these are associated with a wide range of different effectors, such as the transmembrane protein E-cadherin (E-Cad), and together they build up the adherens junction (AJ) [5]. AJs play the role of a mechanical barrier in the tissue, ensuring that cells are closely packed and resistant to physical stress [6].

Basal to AJs, in *Drosophila* epithelia, a second type of junction named septate junction (SJ) – the functional equivalent of tight junctions (TJs) in vertebrates [7] – is involved in the formation of the paracellular diffusion barrier. SJs consist of a large protein complex comprising more than 25 proteins [8, 9]. Among them are cytoplasmic proteins such as Coracle (Cora), glycophosphatidylinositol (GPI)-anchored proteins, single-pass transmembrane proteins such as Neurexin-IV (Nrx-IV), and Claudin-like proteins. At the meeting point of three cells within an epithelium, a specialized domain called a tricellular junction (TCJ) arises, and to date three proteins have been described as enriched at the SJ level: Gliotactin (Gli) [10], Anakonda (Aka; also known as Bark Beetle [11, 12]) and the myelin proteolipid protein family member M6 [13]. We and others have recently described an intricate interplay in which both Aka and M6 are required to recruit and stabilize themselves at the TCJ, while Gli is needed to stabilize them both at the TCJ [14, 15]. Moreover, we have shown that TCJ proteins are required to ensure the anchoring of SJ proteins at the vertex, and, in turn, vertex-specific enrichment and restriction of TCJ proteins are linked to SJ integrity [14].

As described above for AJs, SJs must also be highly plastic to cope with a high rate of cell division, tissue growth, cell intercalation or delamination, while maintaining the integrity of the permeability barrier. Our previous work contributed to show that SJs are stable complexes, exhibiting a turnover rate of 90 minutes. SJ components are delivered and assembled apically, just basal to AJs, and continue to be progressively dragged basally in a treadmill-like manner [16]. At the basal SJ belt, SJ components are thought to be disassembled, internalized and recycled apically to form new SJs or to be degraded. Several studies have revealed that intracellular trafficking actors, such as Rab11 [17], the retromer, and the endosomal sorting complexes required for transport (ESCRT)-III component Shrub, are key regulators of SJ establishment and integrity [18]. Retromer is implicated in the retrieval of cargos from endosomes while ESCRT-III regulates ubiquitin-dependent degradation of transmembrane cargoes. In addition, the Ly6-like proteins Crooked, Coiled, Crimpled [19] and Boudin [20, 21], components of the SJ, have been reported to regulate the endocytic trafficking of Nrx- IV and Claudin-like Kune-Kune, indicating that the SJ components regulate each other’s presence in the SJ.

Despite the fact that SJs have been extensively studied for the past decades, it only recently emerged that they might be involved in additional mechanisms beyond their initially described filtering actions [9]. For instance, a striking feature of *Drosophila* embryo SJ mutants is the appearance of a wavy trachea associated with loss of the permeability barrier. Other morphogenetic defects include diminished and deformed salivary glands, head involution and dorsal closure defects. SJ proteins also regulate the rate of division of intestinal stem cells [22, 23], as well as hemocyte lineage differentiation *via* interactions with the Hippo pathway [24, 25]. Another intriguing feature is the confirmation of the role of SJ components in wound healing [26]. Indeed, lack of different SJ components impairs the formation of actomyosin cables, which are regulated by AJs and under normal conditions ensure the proper healing of the tissue. Hence, the studies cited revealed that SJ proteins can impact mechanical properties of the tissue, calling for a deeper understanding of the impact that the loss of SJ integrity has on general mature tissue homeostasis.

We recently reported that defects at tricellular Septate Junctions (tSJs) are always accompanied by bicellular Septate Junctions (bSJs) defects. Indeed, restriction of tSJ components at the vertex is dependent on bSJ integrity. Conversely, loss of tSJ components causes considerable membrane deformation and the loss of bSJs abutting the vertex [14]. However, and surprisingly, under these conditions, cells remain within the epithelial layer and do not delaminate. In this paper, we investigate how cell adhesion is modulated and allows epithelial integrity to be maintained following disruption of the paracellular diffusion barrier. We use the *Drosophila* pupal notum as a model of mature epithelium with established and functional mechanical and paracellular diffusion barrier functions, a tissue that lends itself to quantitative imaging and in which we can easily dissect the mechanics and genetics of epithelia.

## Results

### Disruption of tri- and bicellular septate junction integrity alters the distribution of adherens junction components

We have previously described that NrxIV-labelled bSJs no longer terminate at vertices when TCJ components are lost [14]. Here, we report that, at the electron microscopy resolution, depletion of Aka induces weaknesses in the integrity of the tissue which results in cell membrane detachment at the vertex in the plane of the SJ and formation of sizeable intercellular gaps within the epithelium (Figure 1A–A′). To know how these gaps impact the overall epithelial integrity, we investigated the relationship between tSJs and AJs. We measured a 2 fold enrichment of *Drosophila* E-Cad tagged with GFP (E-Cad::GFP) at tAJs and 1.5 fold enrichment at bAJs in *aka^L200^* mutant cells (Figure 1B and C). This enrichment was also observed using an E-Cad antibody (data not shown). The E-Cad::GFP increase was accompanied by an enrichment of junctional Myo-II tagged with GFP (Myo-II::GFP) both at tAJs (1.7 fold enrichment) and bAJs (1.8 fold enrichment; Figure 1D and E). The junctional and medial pools of Myo- II act in synergy with forces exerted by the medial-apical meshwork transmitted onto the junctional pool [27]. The medial-apical network was also stronger in *aka^L200^* cells than in wild-type (WT) cells (1.5 fold enrichment; Figure 1D and E). In addition, we probed F-actin and determined that loss of Aka resulted in a 1.9 fold and 2.5 fold increase in staining at bAJs and tAJs, respectively (Figure 1F and G). We observed similar results upon loss of Gli and M6, suggesting that loss of tSJs is responsible for the observed defects. Next, using a hypomorphic allele of the transmembrane bSJ protein Nervana 2 (Nrv2), we found that E-Cad::GFP (Figure S1A and B) and Myo- II::GFP (Figure S1C and D) were enriched at both bAJs and tAJs in *nrv2^k13315^*cells compared with WT cells. E-Cad enrichment was also observed upon loss of GPI-anchored bSJ proteins such as Coiled and Crooked (data not shown), indicating that alteration of the SJ resulted in increased levels of E-Cad in the plane of AJ and thus raises the possibility of concomitant changes in epithelial cell adhesive properties, which we have subsequently studied.

**Figure 1:**
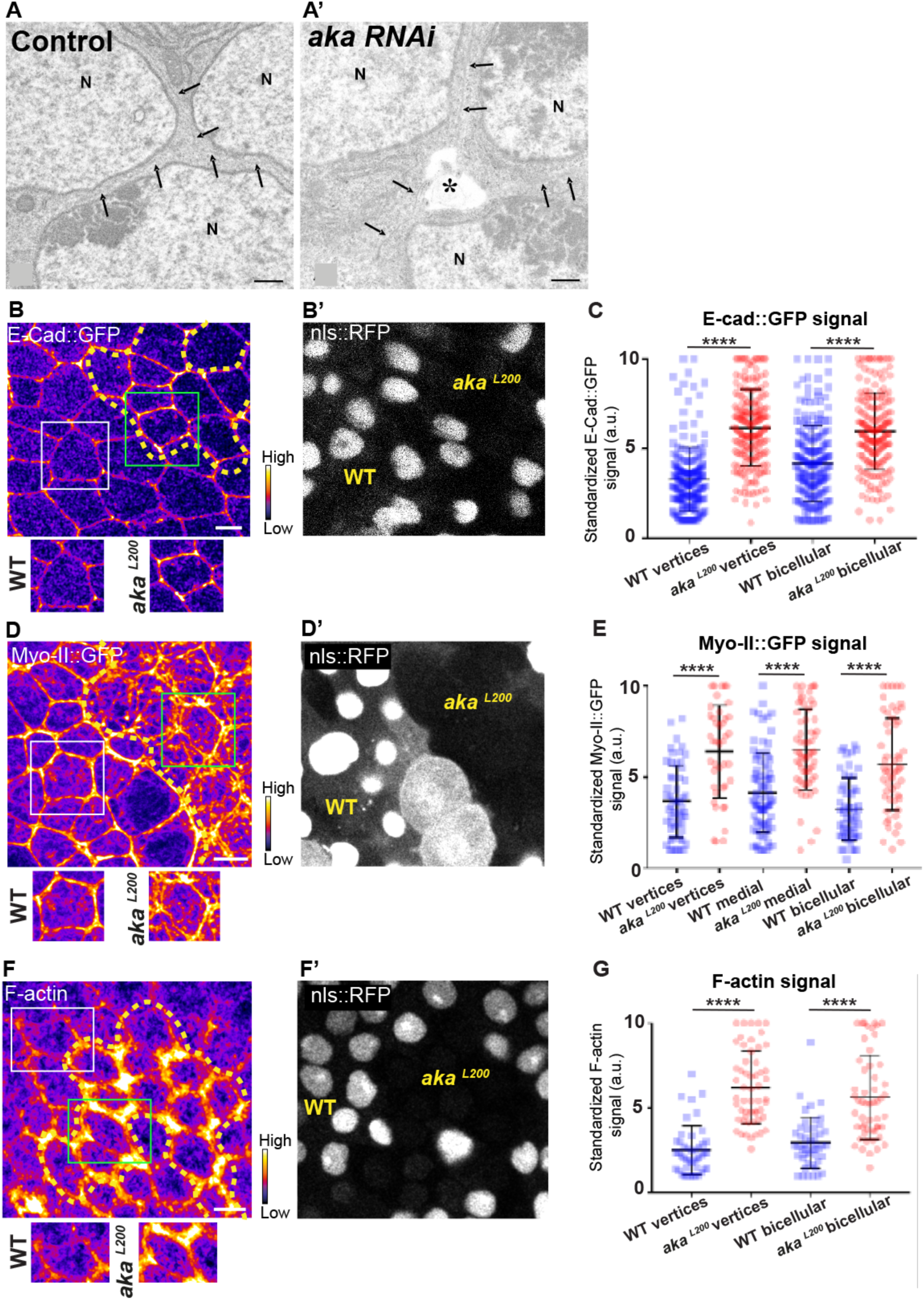
Consequence of loss of Anakonda on tricellular septate junction morphology and adherens junction components. Transmission electron microscopy of wild-type (A) and aka RNAi (A′) pupal notum. Note that Aka-depleted cells present an intercellular gap (asterisk) at the tricellular junction at the level of the nucleus. N: cell nucleus; arrows: cell membranes. (B–B′, D– D′ and F–F′) Localization of E-Cad::GFP (B, fire colour), Myo-II::GFP (D, fire colour) or F-actin (F, phalloidin, fire colour) in wild-type and *aka^L200^* cells. Wild-type cells, identified by nls::RFP (grey in panels B′, D′ and F′), are separated from *aka^L200^* cells by the dashed yellow line. (C) Plot of the standardized E-Cad::GFP signal at bicellular junctions and vertices in wild-type (blue squares) and *aka^L200^* cells (red circles) (n = 201 and 193 vertices and n = 208 and 188 bicellular junctions for wild-type and *aka^L200^* respectively; >5 pupae for each condition). (E) Plot of the standardized Myo-II::GFP signal at bicellular junctions, vertices as well as medial network in wild-type (blue squares) and *aka^L200^* cells (red circles) (n = 54 and 42 vertices and n = 84 and 61 cellular medial networks and n = 55 and 56 bicellular junctions for wild-type and *aka^L200^*, respectively; n = 5 pupae for each condition). (G) Plot of the standardized F- actin signal at bicellular junctions and vertices in wild-type (blue squares) and *aka^L200^* cells (red circles) (n = 45 and 55 vertices and n = 47 and 54 bicellular junctions for wild-type and *aka^L200^*, respectively; n = 5 pupae for each condition). Bars show mean ± SD, ****p < 0.0001, Mann–Whitney test. A calibration bar shows LUT for grey value range. The scale bars represent 500 nm for panels A–A′ and 5 µm for panels B–F′. White squares represent close-up of WT and green squares of *aka^L200^* situations for panels B, D and F.

### The loss of Anakonda alters the adhesive and the mechanical epithelial properties

Because AJs are sites of mechanical force transduction, we hypothesized that the higher levels of E-Cad and Myo-II modify the mechanical properties of the tissue. To assess it, we first tested if Myo-II was activated in *aka* mutant context, by using an antibody against phosphorylated Myo-II (p-Myo-II), and we observed an enrichment in *aka^L200^* cells compared with WT cells (Figure 2A-B). The enrichment was of 1.6 fold at tAJs and bAJs and of 1.8 fold at the medial-apical network (Figure 2A-B). Next, we probed junctional tension using two-photon laser-based nanoablation in the plane of the AJ labelled with E-Cad::GFP (Figure S1E-F). Intriguingly, no significant differences in recoil velocities were observed upon ablation of WT cells versus *aka^L200^* mutant junctions (mean = 0.19 ± 0.08 µm/s in WT vs mean = 0.20 ± 0.07 µm/s in *aka^L200^*) or *nrv2^k13315^* cells (mean = 0.15 ± 0.07 µm/s in WT vs mean = 0.16 ± 0.08 µm/s in *nrv2^k13315^*) (Figure S1F). While recoil velocities indicated that there was no change in in-plane membrane tension upon loss of Aka, we noticed that the cell area of *aka^L200^* cells was slightly reduced by 12% compared to WT (Figure S1G). This prompted us to analyse the length of the new adhesive interface formed during cell cytokinesis. Indeed, when a cell divides and forms its new cell–cell adhesive interface at the AJ level, the length of the new junction is determined by various factors: the force balance between the cells’ autonomous strength in the actomyosin contractile ring, the cells’ non-autonomous response of neighbouring cells that recruit contractile Myo-II at the edges to impose the geometry/length of the new interface, and the strength of intercellular adhesion defining the threshold of disengagement [28–30]. Notably, E- Cad overexpression was reported to delay junction disengagement leading to a shorter interface in early embryos [29]. First, we observed that when a WT cell divides between one WT and one *aka^L200^* cell, the Myo-II::GFP signal was higher during the formation of the new vertex and at the future vertex formed at the interface between WT and *aka^L200^* cell, where there is no Aka (Figure 2C-D; white arrow) compared to the WT interface (Figure 2C-D; green arrow). While this phenomenon can be observed in WT conditions, the proportion of asymmetric enrichment of Myo-II::GFP was much higher in *aka^L200^* conditions (Figure 2E). Then, we confirmed that WT cells established a long E-Cad adhesive interface upon completion of cytokinesis, with few fluctuations in length and across time over the 30 minutes after the onset of anaphase (Figure 2F and H) as expected from [28, 30]. In contrast, *aka^L200^* cells showed a reduction in this junctional length, as highlighted in some extreme cases of shrinkage (Figure 2G and H). This change in the new cell–cell interface length observed in *aka^L200^*cells started to be significant approximately 10 minutes after the onset of anaphase (Figure 2H), suggesting fewer resisting forces from neighbours and/or increased constriction from the dividing cell.

**Figure 2:**
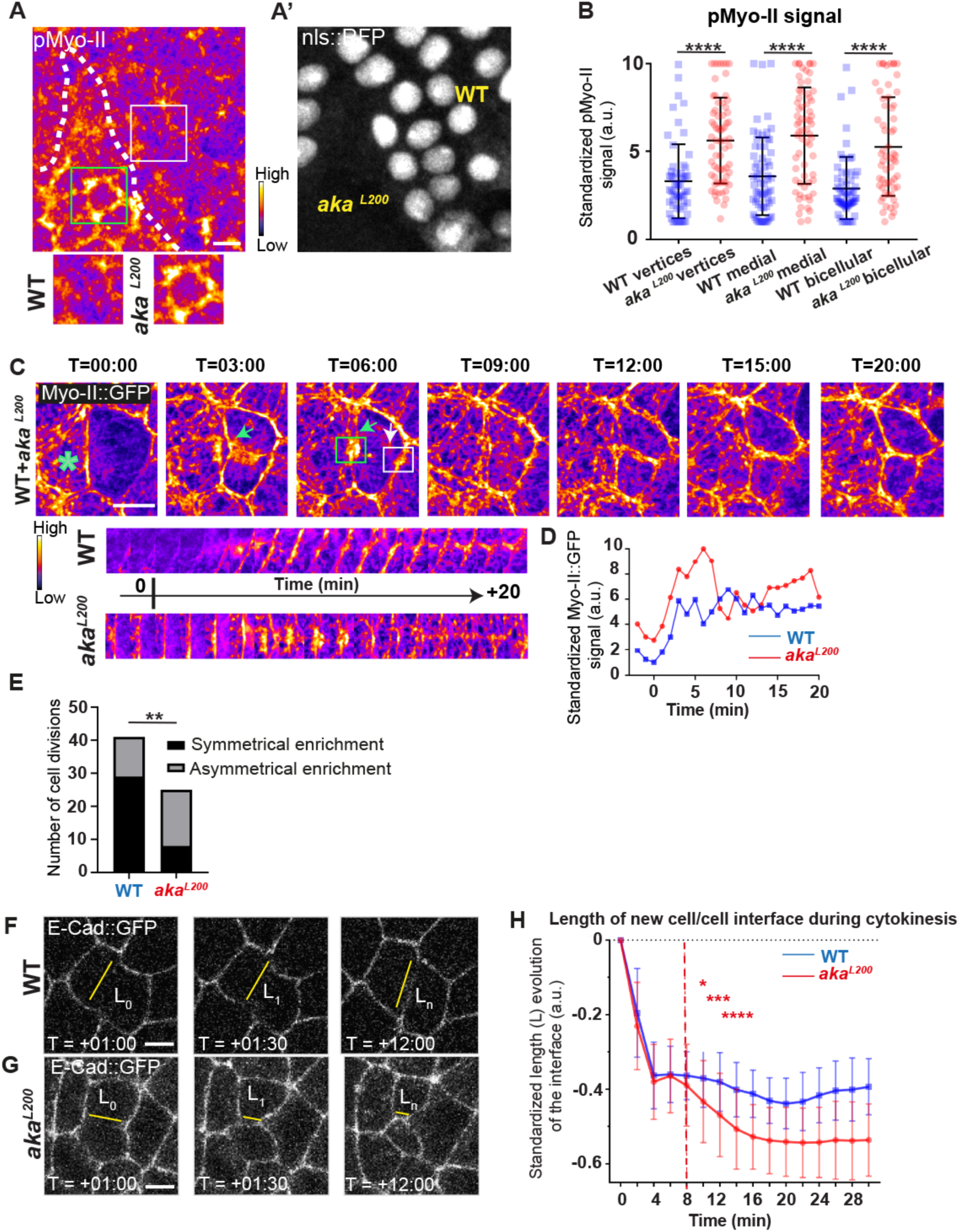
Loss of Anakonda promotes Myo-II activation and mechanical disturbances at adherens junction level during interphase and cytokinesis. (A) Localization of phospho-Myo-II (pMyo-II; fire color) in wild type (nls::RFP positive) and *aka^L200^* mutant cells (nls::RFP negative). (B) Plot of the standardized pMyo-II signal at tri- and bicellular junctions as well as medial network in wild-type (blue squares) and *aka^L200^*cells (red circles) (n = 57 and 67 vertices, n = 65 and 66 cellular medial networks and n = 62 and 61 bicellular junctions for wild-type and *aka^L200^*, respectively; n > 5 pupae for each condition). (C) Cytokinesis of *a WT cell* expressing Myo-II::GFP between 16h-19h APF, after heat-shock to induce clone of wild type and mutant cells for *akaL200*. Representation of a WT cell cytokinesis with recruitment of a higher amount of Myo-II::GFP at the contact with *akaL200* cell (marked by the green asterisk, green arrow for Myo-II::GFP signal) compared to the WT one (white arrow). Myo-II::GFP recruitment is asymmetrical in terms of Myo-II::GFP signal intensity. Kymograph represents the asymmetric enrichment of Myo-II::GFP of the WT and *aka^L200^*newly formed vertices depicted above. (D) Plot representing the Myo-II::GFP signal during cytokinesis at the WT (blue line) and *aka^L200^* (red line) newly formed vertices depicted in C. Time is min:sec with t=0 corresponding to the anaphase onset. (E) Histogram representing the percentage of cells displaying symmetrical (black) or asymmetrical (dark grey) Myo-II::GFP recruitment during cytokinesis of WT with WT neighbours and of WT with one WT and one *aka^L200^* neighbours (n = 29 and n =12 ; n = 8 and n = 17 for symmetrical and asymmetrical enrichment in WT and *aka^L200^* conditions respectively; n > 5 pupae for each condition). (F-H) Cytokinesis of *notum* cells expressing E-Cad::GFP at 16h APF, after heat-shock to induce clone of wild type (F) and mutant cells for *aka^L200^*(G). Time is min:sec with t=0 corresponding to the anaphase onset. L represents the length of the new cell/cell interface. (H) Plot of the mean length interface at each corresponding time points. Wild type situation is represented by blue squares and *aka^L200^* situation is represented by red circles. Bars show Mean ± SD, * p < 0.05, ** p < 0.005, *** p = 0.0001, **** p < 0.0001, unpaired t test and Mann-Whitney test for panels B, Fisher t test for panel E and Multiple t test for panel H. A calibration bar shows LUT for grey value range. The scale bars represent 5µm. White square represents close-up of WT and green square of *aka^L200^* situations for panel A.

To further explore defects in adhesive properties and mechanical tension caused upon loss of Aka, we examined the localization of Vinculin (Vinc), an F-actin binding partner recruited at junctions in a tension-dependent manner [31, 32]. We observed higher levels of GFP-tagged Vinc (Vinc::GFP) at tAJs (2 fold enrichment) and bAJs (1.75 fold enrichment) in *aka^L200^* cells compared with WT cells (Figure S2A and B). Strikingly, upon loss of Aka, Vinc::GFP was found enriched not only at the AJ level but also at the basal part of mutant cells (Figure S2C–D′; see below), raising the possibility of a reorganization of the F-actin-anchoring point to the membrane associated with increased tension at these localizations (see below). We also found that another F-actin crosslinker, Karst, was enriched at the AJ level at bAJs (1.4 fold enrichment), at tAJs (60% enrichment) and at the apical–medial part of the cell (1.2 fold enrichment; Figure S2E and F). Then, we investigated the localization of the Hippo/YAP partner Ajuba (Jub), known to be increased at AJ upon increased tension in *Drosophila* wing discs [33]. We observed an increase of GFP-tagged Jub (Jub::GFP) marking at tAJs (1.4 fold enrichment) and at bAJs (1.75 fold enrichment) (Figure S2G-H). Collectively, these results suggest that the loss of Aka and concomitant disruption of SJ integrity increase apical tension and/or adhesive properties in epithelial cells. The mechanisms through which alteration of SJ components impact AJ were then investigated.

### Septate junction alterations are associated with ESCRT complex defects

Several studies have revealed that the establishment and integrity of bSJs rely on intracellular traffic [18, 19, 34]. Among them, Vps35 subcellular localization is regulated by Shrub, which is itself needed to ensure correct bSJ protein delivery at the plasma membrane. In the pupal epithelium, loss of Shrub causes loss of ATP-α::GFP signal, indicative of an interplay between SJs and endosomal sorting machinery [18].

Upon loss of Aka, bSJs are no longer connected to vertices and exhibit membrane deformation with increased levels of bSJ components [14]. The higher level of bSJ components could result from an increased delivery of newly synthesized proteins, reduced endocytosis, and/or increased recycling of bSJ proteins. We hypothesize that defects in SJ integrity might feedback on the endocytosis-recycling of bSJ proteins, to compensate for SJ defects. To probe for possible membrane traffic alterations, we investigated the ESCRT complex by examining the multivesicular body (MVB) marker, the ESCRT-0 component hepatocyte-growth-factor-regulated tyrosine kinase substrate (HRS)/Vps27 and Shrub/Vps32 endogenously tagged with GFP (Shrub::GFP). In the control situation (Figure 3A–B′), HRS and Shrub::GFP appeared as small punctate structures that substantially colocalized (white structures; Figure 3A–B′). Strikingly, silencing of the core bSJ protein Cora induced the formation of enlarged Shrub::GFP-positive structures, more and larger HRS positive compartments (Figure 3C–D′), together with bSJ integrity alteration (Figure 3C’). The enlarged Shrub::GFP-positive structures did not colocalize with HRS punctae (Figure 3C–D′). We also detected larger and brighter HRS-positive structures, both in *aka^L200^*(Figure S3A and B) and *nrv2^k13315^* cells (Figure S3C and D).

**Figure 3:**
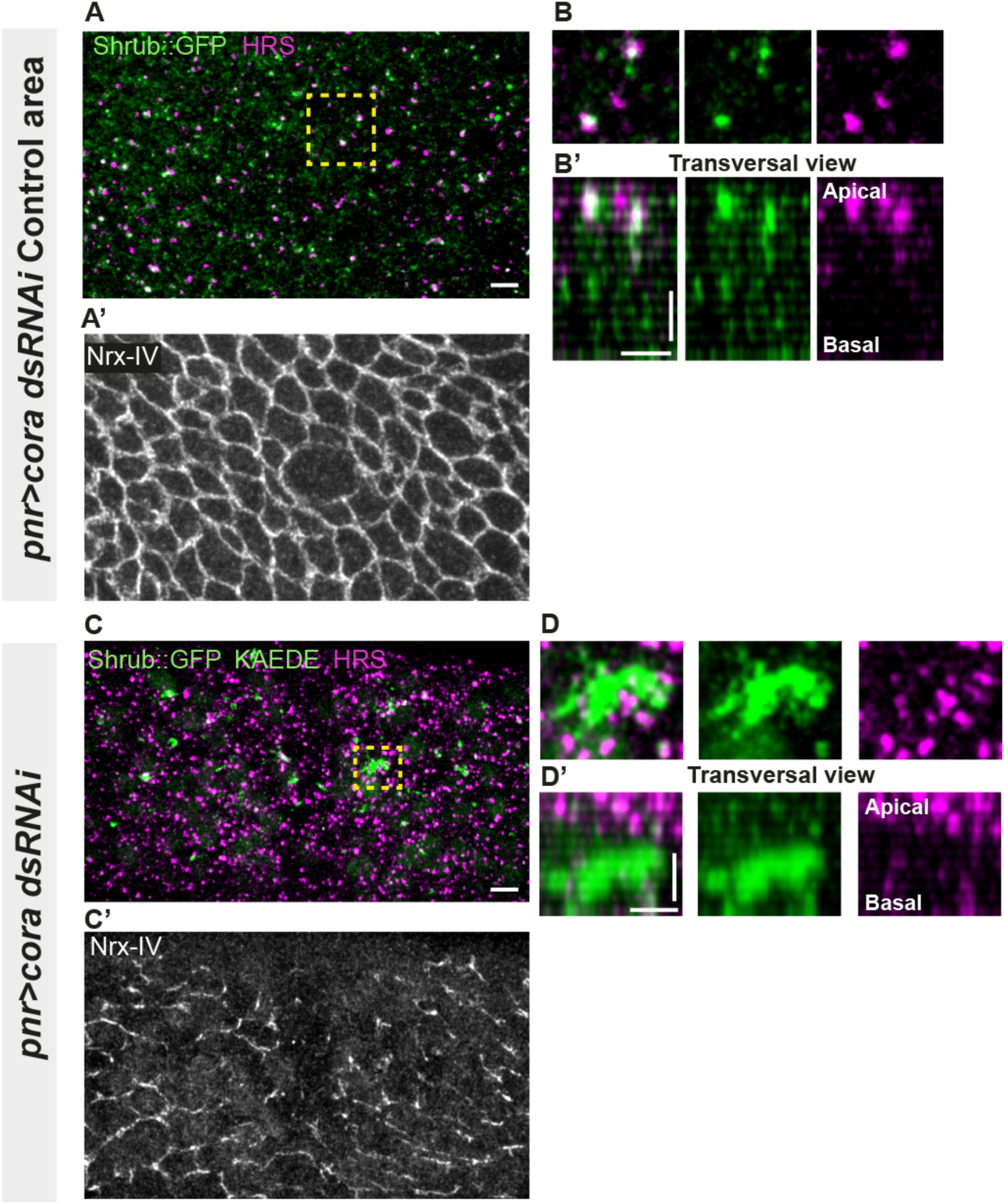
Septate junction defects are associated with increased number of HRS- and ESCRT III protein Shrub-positive structures. (A-B’ and C-D’) Localization of Shrub::GFP + GFP antibody (green), KAEDE (A-F’) in cells marked by Nrx-IV (anti-Nrx-IV, grey) and HRS (anti-HRS, magenta) in wild type and cells expressing UAS::*cora*-RNAi together with UAS::KAEDE under *pnr*-Gal4 control. (A-B’) Localization of Shrub::GFP + GFP antibody and HRS in a wild-type area of a tissue expressing UAS::*cora*-RNAi and UAS::KAEDE under *pnr*-Gal4 control (KAEDE negative and regular Nrx-IV signal in (A’) in a planar view (A, A’ and B) or in a transversal view (B’). Yellow dashed square shows (B and B’) magnification of wild type cell with colocalization between Shrub::GFP and HRS at SJ level showed by Nrx-IV. (C-D’) Localization of Shrub::GFP + GFP antibody and HRS in cells expressing UAS::*cora*-RNAi and UAS::KAEDE under *pnr*-Gal4 control (KAEDE positives and Nrx- IV reduced signal in (C’)) in a planar view (C, C’ and D) or in a transversal view (D’). Yellow dashed squares show (D-D’) magnification of aggregates of Shrub::GFP surrounded by HRS staining. The scale bar represents 5µm (A and C) and 3µm in (B’ and D’).

Because the ESCRT complex is involved in controlling the degradation of poly- ubiquitinylated cargoes [35], we then asked whether the excess of Shrub-positive, enlarged structures was due to a change in Shrub degradation activity. A way to probe putative defects in ESCRT function is to monitor the amount of poly-ubiquitinylated proteins targeted for degradation [35]. First, we used an RNAi against Shrub and confirmed that depletion of Shrub led to both poly-ubiquitinylated proteins aggregates appearance, marked using anti-FK2, and SJ alterations as observed by the inhomogeneous Nrx-IV signal (Figure 4A). Then, we studied SJ disruption case using Cora-RNAi approach. In the control situation, Shrub::GFP and poly-ubiquitinylated proteins FK2 formed small punctate compartments (Figure 4B). In striking contrast, in the Cora-depleted domain, Shrub::GFP and anti-FK2 labelled large structures (Figure 4C–C′′). Shrub::GFP-positive structures were closely juxtaposed and/or partially colocalized with FK2, (Figure 4C′–C′′). Similar observations were made upon knock- down of Nrx-IV (Figure 4D–D′′), as well as in *aka^L200^* cells (Figure 4E). Hence, mutants with disrupted SJ integrity display features of a dysfunctional ESCRT-III-dependent degradation pathway, somewhat reminiscent of a *shrub* loss-of-function. Despite these apparent similarities, we noticed that, in contrast to Shrub depletion [36], NrxIV did not accumulate in enlarged intracellular compartments upon Cora depletion. In other words, the accumulation of Shrub::GFP in enlarged compartments seen upon Cora depletion is not functionally equivalent to the loss of Shrub. We propose that it is the Shrub activity that is being modified upon SJ alteration, preventing SJ component degradation in favour of SJ component recycling. In support of this proposal of increased recycling, loss of TCJ components was shown to cause membrane deformations enriched in SJ components [14]. The next question was whether deregulation of Shrub activity by SJ component depletion could affect adhesive properties and cell mechanics.

**Figure 4:**
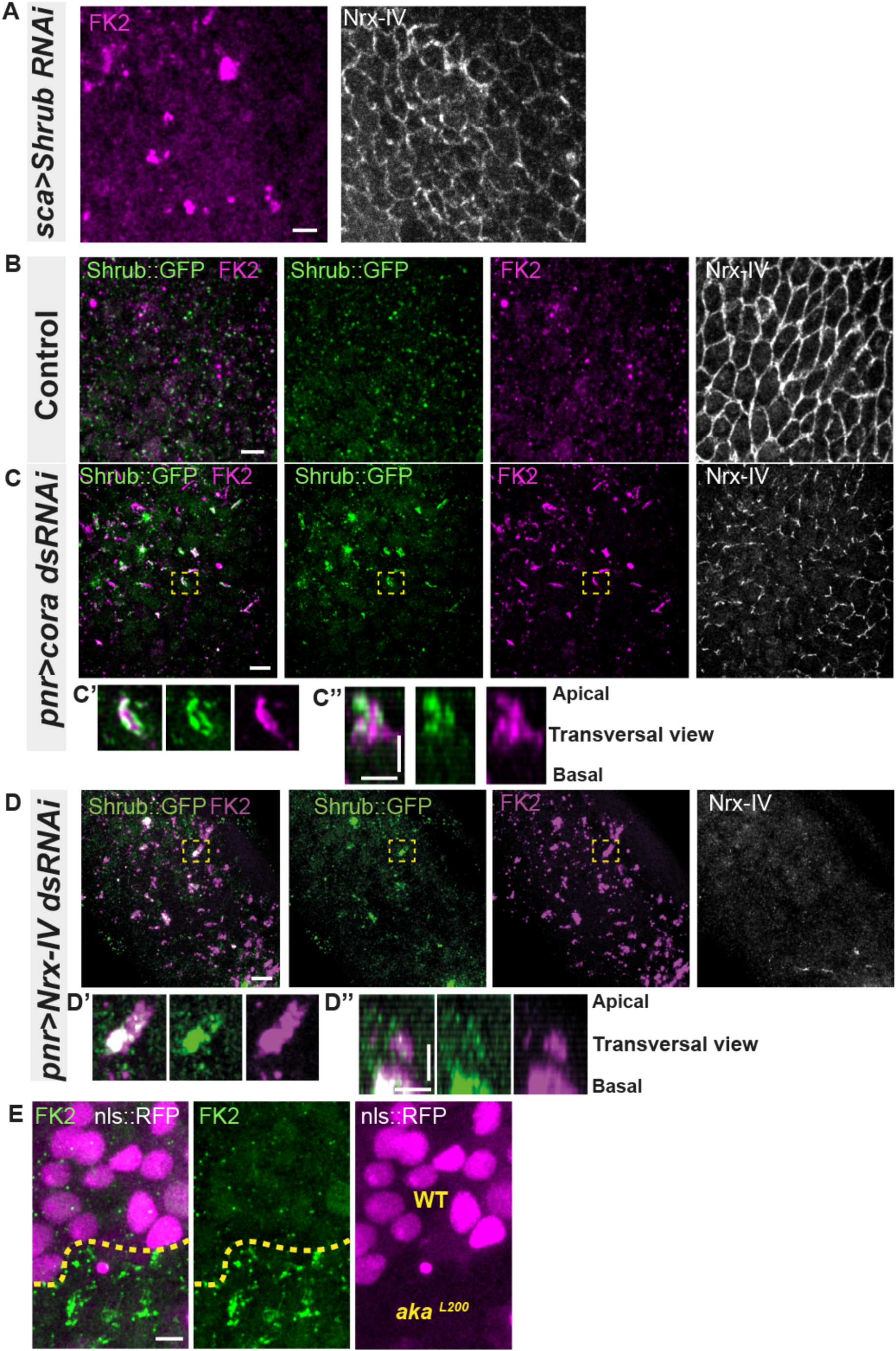
Septate junction defects leads to the enlargement of ESCRT III protein Shrub- and ubiquitinylated proteins- positives structures. (A) Localization of FK2 (anti ubiquitinylated proteins, magenta) in cells marked by Nrx- IV (anti-Nrx-IV, white) expressing UAS::*shrub*-RNAi under *sca*-Gal4 control. (B-C’) Localization of Shrub::GFP + GFP antibody (green), in cells marked by Nrx-IV (anti- Nrx-IV, white) and FK2 (anti ubiquitinylated proteins, magenta) in wild type and cells expressing UAS::*cora*-RNAi or UAS::*Nrx-IV*-RNAi under *pnr*-Gal4 control. (B-C’’) Localization of Shrb::GFP + GFP antibody and FK2 in a wild-type area (regular Nrx- IV signal in (B)) or in cells expressing UAS::*cora*-RNAi under *pnr*-Gal4 control (Nrx-IV reduced signal in (C)). Yellow dashed square shows (C’ and C’’) magnification of cells with partial or no colocalization between Shrb::GFP and FK2 as well as aggregates of FK2 surrounded by Shrb::GFP staining in a planar view (C’) and transversal view (C’’). (D-D’’) Localization of Shrub::GFP + GFP antibody and FK2 in cells expressing UAS::*Nrx-IV*-RNAi under *pnr*-Gal4 control (Nrx-IV signal disappearance in (D)). Yellow dashed square shows (D’ and D’’) magnification of cells with partial or no colocalization between Shrb::GFP and FK2 in a planar view (D’) and transversal view (D’’). (E) Localization of FK2 in both wild-type and *aka^L200^* cells, separated by the dashed yellow line. Clones of wild-type and *aka^L200^*cells identified by nls::RFP marking (magenta). The scale bar represents 5µm (A, B, C, D and E) and 3µm in (C’’ and D’’).

### Loss of tricellular or bicellular septate junction components impact Crumbs localization and triggers assembly of focal adhesion contacts

In *Drosophila* trachea, loss of Shrub has been reported to affect the localization of bSJ components, such as Kune-Kune, impairing the paracellular diffusion barrier on one hand and Crb activity on the other [17]. Loss of Shrub results in an elongated sinusoidal tube phenotype which was shown to be caused by mislocalized Crb activation. Indeed, in *shrb^4^* clones, instead of being restricted to the junctional domain, Crb is present in ESCRT-0-positive endosomal compartments causing Crb overactivation [17]. In this study, the authors raised the possibility that the defect of bSJ caused by loss of Shrub might also contribute to Crb activation, a possibility that we tested next. As a control, we monitored the localization of SJ protein Kune Kune (Kune) and Crb using an anti-Crb antibody targeting its N-terminal extracellular domain (anti-Crb). We observed a colocalization in small vesicles at the basal level of the cell (white vesicles; Figure S4A-A’’), suggesting that Kune and Crb traffic together. Upon knock-down of Shrub via RNAi, we observed defects of Kune and Crb characterized by enrichment of Crb and Kune in basal aggregates (Figure S4B-B’’’). The apparent similarities between depletion of Shrub and that of b/tSJ components on FK2 and HRS, raised the question whether the loss of Aka could result in defective Crumbs localization, that we next investigated. We monitored Crb localization in tSJ defects situation using Crb tagged with a GFP in its extracellular domain (Crb::GFP) or an anti-Crb antibody in *aka^L200^* context. Crb signal was detected both at junctional and medial apical parts of WT cells (Figure 5A and C). Strikingly, in *aka^L200^* and in bSJ defective *nrv2^k13315^* cells, the apical–medial Crb signal was increased (Figure 5A–C′ and Figure S5A–C). Concerning junctional Crb, we observed both an enrichment associated with small aggregated structures at or adjacent to the junctions using Crb::GFP in *aka^L200^* cells (Figure 5A–A′′) and a less well-defined signal compared to Crb::GFP when using the anti-Crb (Figure 5B–B′′). In *nrv2^k13315^*cells, although junctional Crb::GFP signal was not significantly different than in control cells, anti-Crb antibody showed a less well-defined signal compared to Crb::GFP at the junction, suggesting that, perhaps, Crumbs is closely juxtaposed to the plasma membrane rather than residing at the plasma membrane (Figure S5A–C). Interestingly and in striking contrast to Shrub depletion, we did not observe Crb and Kune basal aggregates in *aka^L200^* and *nrv2^k13315^* conditions (data not shown). Hence, if both Shrub and bSJ/tSJ defects lead to Crumb enhanced signals, Shrub depletion is responsible for Crb being enriched in enlarged compartments whereas loss of Aka or Nrv2 triggers Crb enrichment at the apical level of the cell. Thus, as proposed above for Nrx-IV, these data further suggest a hijacking of Shrub activity toward recycling components upon alteration of SJ integrity. The elevated apical levels of Crb upon depletion of SJ component is proposed to be causal to apical enrichment of the Crumbs effector Karst (Figure S2E and F) [37]. Therefore, we decided to remove one copy of Crb in the *aka^L200^* context to observe if we were able to rescue the AJ phenotype. Although we observed a rescue of the cell area phenotype (Figure 5F), removal of one copy of Crb was not sufficient to restore E- Cad::GFP level to the control situation (Figure 5D-E, 1.7 fold enrichment for bicellular junctions, 1.8 fold enrichment for tricellular junctions).

**Figure 5:**
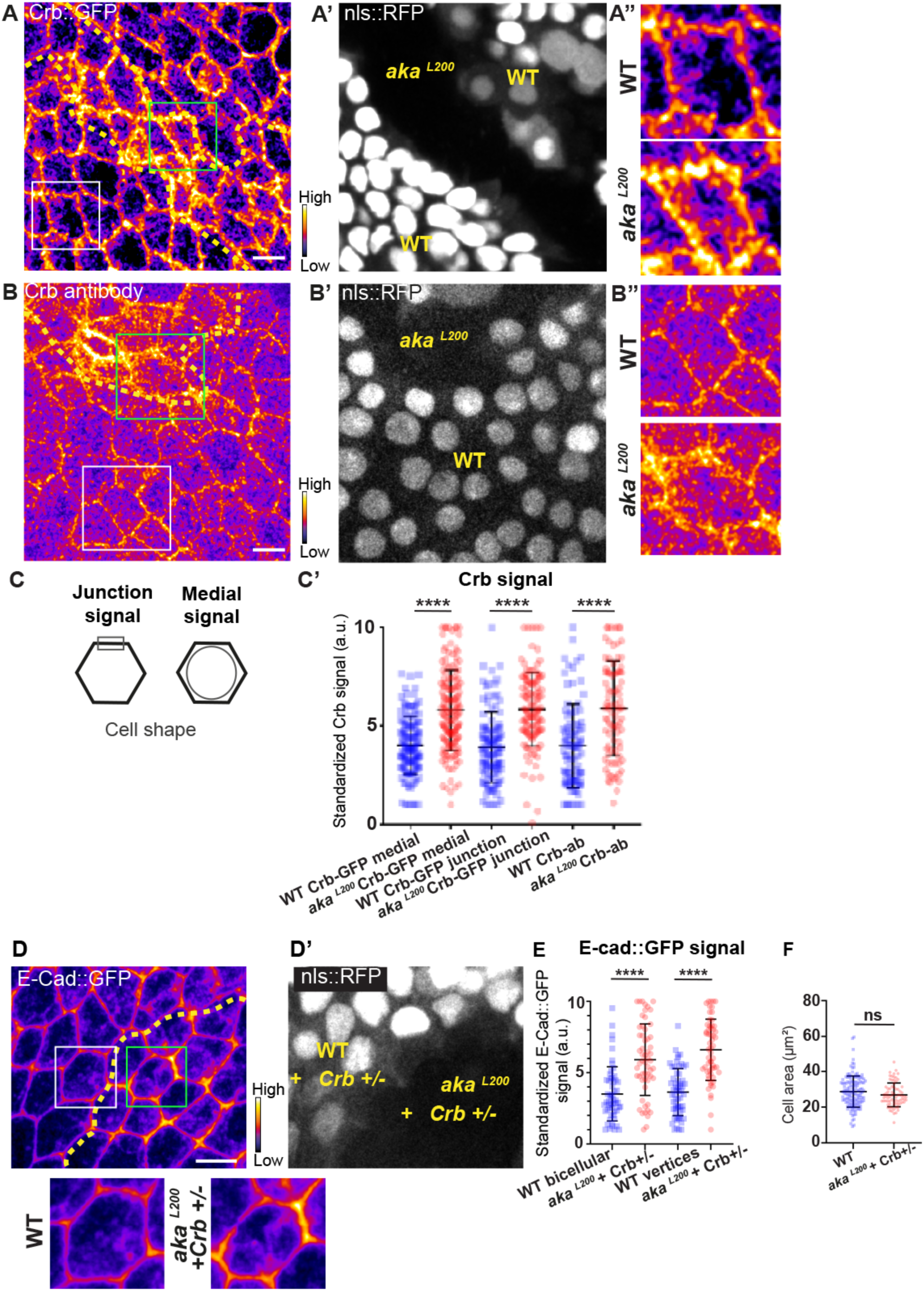
Loss of Anakonda leads to higher level of Crumbs at both junctional and medial part of the cell. (A-A’) Localization of Crb::GFP in both wild-type and *aka^L200^* cells, separated by the dashed yellow line. (A’) Clones of wild-type and *aka^L200^* cells identified by nls::RFP marking. White squares show (A’’) magnification of cell depicted in (A). (B-B’’) Localization of anti-Crb in both wild-type and *aka^L200^* cells, separated by the dashed yellow line. (B’) Clones of wild-type and *aka^L200^*cells identified by nls::RFP marking. White squares show (B’’) magnification of cell depicted in (B). (C) Scheme representing junctional and medial population of Crb staining. (C’) Plot of the standardized Crb::GFP signal at the medial and junctional part of the cell or Crb-ab only at the medial part, in wild type (blue squares) and *aka^L200^* cells (red circles). (n = 100 and 96 cellular medial networks with Crb::GFP, n = 110 and 119 junctions with Crb::GFP and n = 90 and 88 cellular medial networks with Crb-ab for wild-type and *aka^L200^* respectively, n = 5 pupae for each condition). (D-D’) Localization of E- Cad::GFP (D, fire colour) in wild-type (D′, nls::RFP-positive cells in grey) and *aka^L200^* (D′, nls::RFP negative) cells lacking one copy of Crb (Crb+/-). Wild-type and *aka^L200^*cells are separated by the dashed yellow lines in (D–D′). Higher magnification of yellow dashed square depicted in panels D for wild-type and *aka^L200^* cells. (E) Plot of the standardized E-Cad::GFP signal at bicellular junctions and vertices in wild-type (blue squares) and *aka^L200^* (red circles) cells lacking one copy of Crb (n = 55 and 57 bicellular junctions and n = 59 and 58 vertices for wild-type and *aka^L200^* cells respectively; n = 4 pupae for each condition). (F) Quantification of the cell area (in µm^2^) of WT (blue squares, n = 136 cells, n = 4 pupae) and *aka^L200^* cells lacking one copy of Crb (red circles, n = 75 cells, n = 4 pupae). Bars show Mean ± SD, **** p < 0.0001, Mann- Whitney test. A calibration bar shows LUT for grey value range. The scale bars represent 5µm. White squares represent close-up of WT and green squares of *aka^L200^*situations for panels A, B and D.

Loss of Aka leads to elevated Crb, E-Cad, p-Myo-II and Vinc::GFP signals at AJ level. In addition, Vinc-GFP staining also increased basally, with Vinc-GFP-positive structures appearing at the basolateral domain (Figure S2C and D). Vinc is recruited both at AJ and in FA contact [31, 32, 38] and α5- and β1-integrins are regulated via the ESCRT pathway in vertebrates [39]. In pupal notum, depletion of Shrub leads to accumulation of Myospheroid (Mys), the β subunit of Drosophila integrin dimer, in basal compartments that partially colocalize with Kune (Figure S6A-A’’’’), presumably enlarged endosomes, indicating that in invertebrate also, β-integrin levels rely on ESCRT-III function. In line with the hypothesis of the hijacking of Shrub activity upon depletion of SJ components, increased levels of integrin were predicted to recycled back to the plasma membrane.

Indeed, and in striking contrast to Shrub depletion, we found that Mys levels are elevated in *aka^L200^*clones, and that Mys localized in basal clusters along with F-actin (Figure 6A and B). Mys also colocalized with Vinc-GFP in *aka^L200^* cells, indicating an assembly of FA contacts in *aka^L200^* mutant cells (Figures 6C and D). Could these FA contact exert forces in aka*^L200^* cells and hence, mutant cells react by increasing their amount of apical E-Cad, perhaps to sustain cell adhesion and prevent cell extrusion? To investigate this possibility, we knock-downed Mys in *aka^L200^* cells. When compared to *aka^L200^*cells (Figure 1 B-C), depletion of Mys in *aka^L200^* cells almost abolished the E-Cad enrichment at bAJs and at tAJs (Figure 6E-F). The cell area was also no longer significantly different than from WT (Figure 6G). Thus, concomitant loss of tSJ and FA contact in mature epithelium is not sufficient to induce cell extrusion. We propose that disruption of SJ barrier in pupal notum redirects Shrub activity to promote recycling of the junctional components Crumbs and Mys that collectively contribute to the maintenance of epithelial integrity.

**Figure 6:**
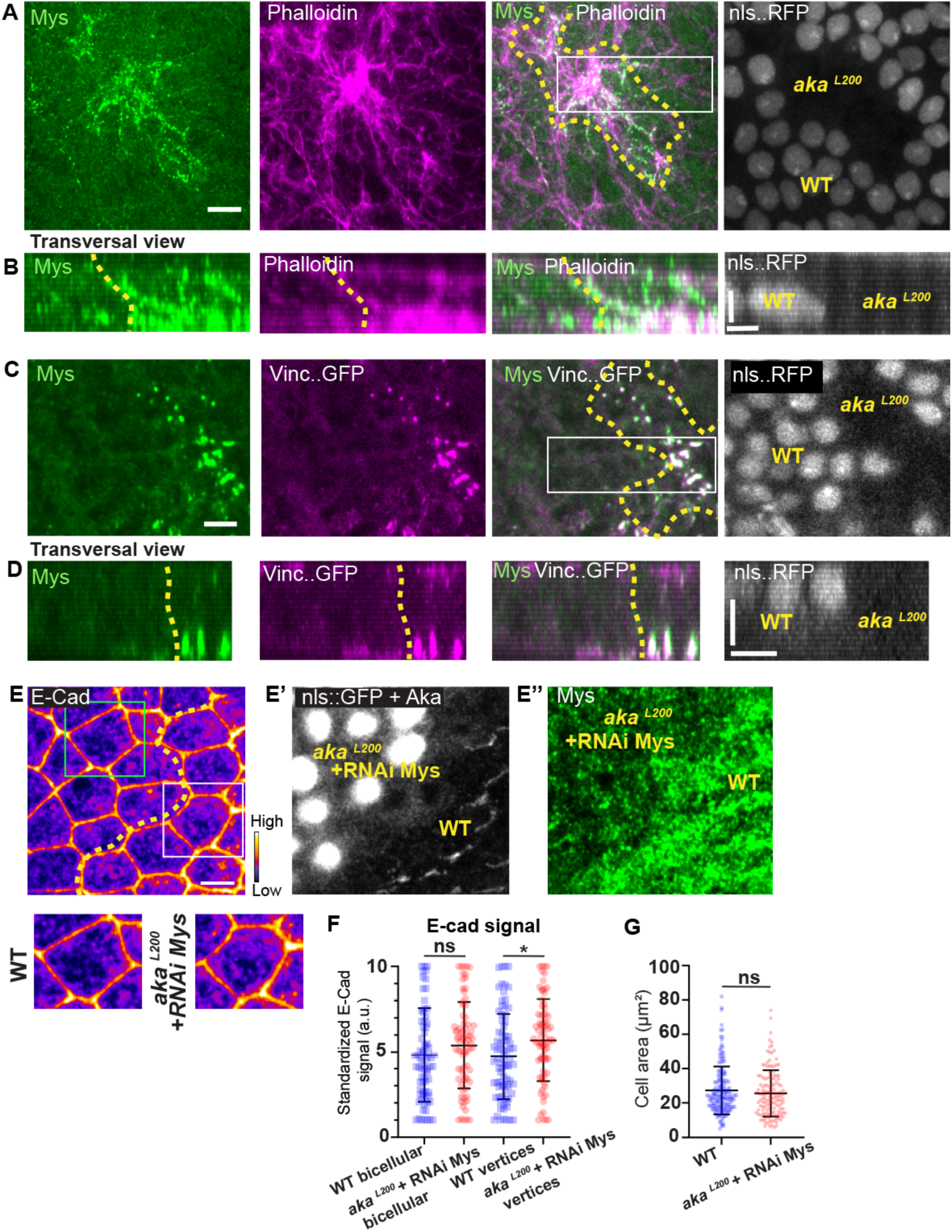
Loss of Anakonda triggers formation of focal adhesions contact. (A) Localization of Mys (green) and F-actin (magenta) in both wild-type and *aka^L200^* cells in a planar view at the basal level, separated by the dashed yellow line. (B) Transversal view of images depicted in A. (C) Localization of Mys (green) and Vinc::GFP (magenta) in both wild-type and *aka^L200^*cells in a planar view at the basal level, separated by the dashed yellow line. (D) Transversal view of images depicted in C. (n > 5 pupae for each condition). (E) Localization of E-Cad (anti-E-Cad; E, fire colour) and Mys stained with Mys antibody (E’’, green color) in wild-type (E′, nls::RFP- negative cells in grey) and *aka^L200^* (D′, nls::RFP positive) cells in which Mys is knock- downed (RNAi-mys). Wild-type and *aka^L200^* cells are separated by the dashed yellow lines in (E). Higher magnification of yellow dashed square depicted in panels E for wild-type and *aka^L200^* cells. (E) Plot of the standardized E-Cad signal at bicellular junctions and vertices in wild-type (blue squares) and *aka^L200^* + Mys knock-downed cells (red circles) (n = 76 and 76 bicellular junctions and n = 81 and 76 vertices for wild-type and *aka^L200^* cells respectively; n > 5 pupae for each condition). (F) Quantification of the cell area (in µm^2^) of WT (blue squares, n = 171 cells, n > 5 pupae) and *aka^L200^* + Mys knock-downed cells (red circles, n = 139 cells, n > 5 pupae). Bars show Mean ± SD, * p < 0.05, Mann-Whitney test. A calibration bar shows LUT for grey value range. The scale bars represent 5µm in A and C and E and 3 µm in B and D. White square represents close-up of WT and green square of *aka^L200^* situations for panel E.

**Figure 7:**
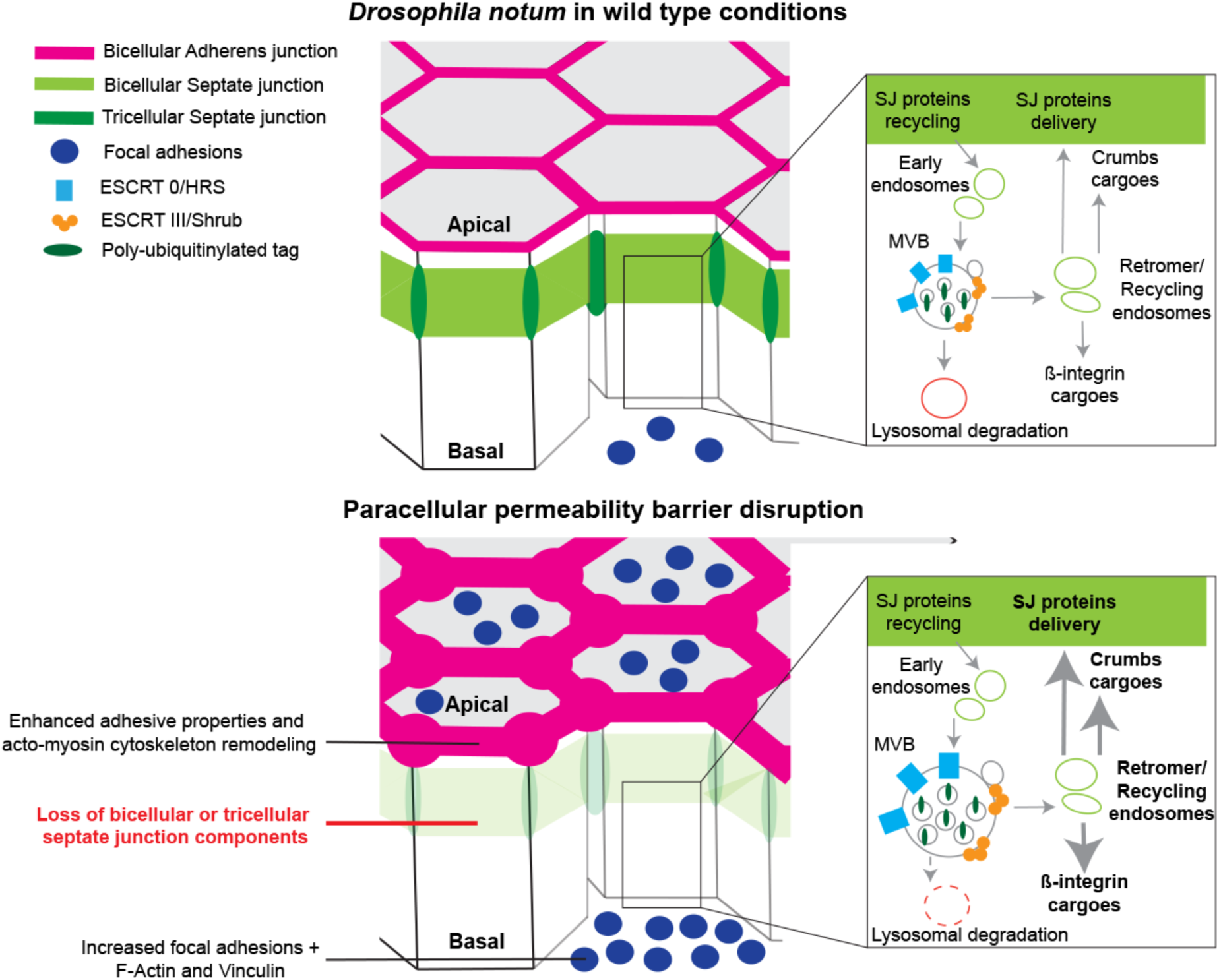
Model summarizing the effects of the disruption of SJ integrity in pupal notum. In wild type conditions, bSJ proteins, β-Integrin and Crumbs are recycled to the membrane thanks to the endosomal-retromer complex. When the paracellular permeability function is compromised due to the loss of SJ components, cells favour recycling over degradation, leading to increased levels of β-Integrin and Crumbs at the cell membrane. The accumulation of β-Integrin and Crumbs leads to a strengthening of the adhesive structure as shown by increased quantity of AJ proteins but also by the appearance of more focal adhesion contacts. We propose that the cell compensates the lack of bSJ contacts by increasing its adhesive properties.

## Discussion

In this study, we examined how epithelial cells can cope with and are able to remain within the tissue upon loss of septate junction integrity. We report that loss of bSJs and tSJs by altering the paracellular diffusion barrier triggers an ESCRT-dependent response to favour bSJ membrane protein recycling instead of promoting lysosomal degradation. By reducing the ESCRT-dependent degradative pathway, the cellular levels of ESCRT cargoes, including Crb and Mys, become elevated. First, we propose that Crb overactivation might be, at least in part, responsible for the change in apical actomyosin contractility/cellular mechanics. Secondly, FA contact points, containing Vinc and Mys, are assembled. We propose a model whereby Crb overactivation and FA contact formation may compensate for bSJ contact alteration, by reinforcing adhesion, ensuring mechanical barrier integrity.

### A mechanism sensing bicellular septate junction defects

In the pupal *notum*, the loss of tSJs leads to a loss of bSJ signal at the vertex [14], weakening the three-cells contact ultimately leading to gaps [this study], preventing cells from fulfilling their role of paracellular diffusion barrier. We propose, based on our previous study [14] and the work of [40] and [41] in which SJ defects have been shown to trigger large membrane deformations, that epithelial cells are capable of detecting SJ defects. Our work shows that a part of the sensing mechanism involves the ESCRT machinery. This machinery exhibits two main functions in endosomal sorting. First, at the outer surface of nascent MVBs, ESCRT machinery is involved in the targeting of ubiquitinylated proteins into intraluminal vesicles, which contain the cargoes destined for lysosomal degradation. Secondly, ESCRT machinery regulates retromer-dependent recycling of bSJ components. The accumulation of the FK2 protein shown here indicates that the primary function of Shrub is attenuated upon alteration of SJ integrity, and we propose that it is in favour of the recycling function. The increased recycling of bSJ components occasioned by the loss of tSJs would contribute to the appearance of large membrane deformations containing an excess of bSJ components. In contrast, the loss of bSJ components, such as Nrv2, Cora or Nrx-IV leads to an overall reduction in the bSJ components Cora, Nrx-IV, ATP-α and Kune-Kune. Indeed, the removal of one of the core components of bSJ prevents assembly and stabilization of bSJs, as shown by fluorescence recovery after photo bleaching (FRAP) analysis [16, 42]. Importantly, it is not the case upon the loss of tSJ proteins [16]. Hence, bSJ-positive membrane deformations appear only upon the loss of tSJ proteins. However, in both cases, the degradative function of Shrub is affected, as attested by the hyperactivation of Crb and the strengthening of FAs (see below).

How can cells sense SJ alteration, modify Shrub function and impact intracellular trafficking? Is it due to the sensing of defects in the paracellular diffusion barrier or in cell adhesive properties, or a combination of both? It is interesting to note that numerous SJ components are GPI-anchored proteins and that, for example, *wunen-1* and *wunen-2* encode lipid phosphate phosphatase [43], so the lipid composition of the lateral plasma membrane is likely to be affected upon the loss of SJ components. In vertebrates, the permeability of the blood–brain barrier is regulated by the major facilitator superfamily domain containing 2a (Mfsd2a) [44]. Mfsd2a is a central nervous system (CNS) endothelial-cell-specific lipid transporter that delivers the omega3-fatty acid docosahexaenoic acid into the brain via transcytosis. Lipids transported by Mfsd2a create a unique lipid composition in CNS endothelial cells that specifically inhibits caveolae-mediated transcytosis to maintain the blood-brain barrier integrity [45]. By analogy to Mfsd2a, in *Drosophila* pupal *notum*, changes in the lipid transported or in the lipid content of the plasma membrane could be sensed upon alteration of SJ integrity and modify intracellular trafficking (i.e. ESCRT-dependent recycling of SJ components). Changes in lateral plasma membrane lipid composition upon SJ alteration could also impact the lipid composition of endosomal compartments that, in turn, could participate in modulating the recycling versus the degradative function of ESCRT [46–48].

### Consequences of septate junction alteration on cellular mechanics and adhesion

In both bSJ and tSJ mutant cells, Crb is enriched at the apical pole of the cells. As Crb is a known binding partner of the β-Heavy Spectrin Karst [37], Crb defects are proposed to cause the enrichment of Karst in the bSJ/tSJ mutant cells. Furthermore, the enrichment of Myo-II::GFP, and especially p-Myo-II, might be due to the upregulation of the activator Rho-kinase (Rok), another known partner of Crb [49]. Moreover, Rok activates Moesin, which links the cell membrane to the actin cortex [50], and Moesin binds to Crb via its FERM domain [51]. Because Moesin is involved in constructing a rigid cortical actin (for example, leading to cell rounding during mitosis [52, 53]), the enrichment of Crb could explain why F-actin and Myo-II enrichment is observed in *aka* mutant cells, even though removing one copy of Crb was not enough to rescue the phenotype, and indirectly explain the change in E-Cad levels and in the tensile forces. Interestingly, the *Drosophila* tSJ protein M6 has been recently reported to act as an interplay partner of Ajuba [54] and loss of M6 is associated with elevated signal of Ajuba at vertices. The fact that we similarly observed elevated signal of Ajuba upon loss of Aka, reinforce the idea of AJ remodelling by direct links between tSJ and AJ/acto-myosin cytoskeleton components. However, we did not observe differences in the recoil velocity of Aka or Nrv2 mutant cells upon laser ablation. A plausible explanation seems related to the fact that all mutant cells have their actomyosin and linked AJs remodelled. Therefore, the pulling forces are homogenous, as in WT conditions, and might be equal on both sides of the junctions. Hence, ablation of a boundary between WT and mutant cells alone might be informative to observe a potential increase in the recoil velocity if we assume from all other data that both populations exhibit an overall difference in force balance. However, this cannot be done because in both the *aka* and *nrv2* contexts, boundaries between WT and mutant cells, display enrichment of E-Cad, Myo-II and F-actin. Another explanation could be that a higher amount of E-Cad and associated actomyosin enhanced the friction at the junctional level, therefore reducing the eventual extra pulling forces [55]. An argument in favour of similar tension in both WT/heterozygous and mutant cells is that clones of mutant cells remained cohesive and not dispersed among WT cells (or vice versa). One could expect mixing of cells upon differential tension at boundaries, as highlighted in [56]. An argument for changes in tensile forces in *aka* mutant cells is the formation of shorter cell–cell interfaces during cytokinesis. However, although these short interfaces could result from high contractile forces within the cytokinetic ring and reduced resistance from neighbours, they could also be the consequence of higher levels of E-Cad and the subsequent delay in E-Cad disengagement, as reported in *Drosophila* embryos [29]. Last but not least, the enhanced adhesion between cells and the extracellular matrix, as revealed by the increase presence of Mys and Vinc, could be a potent explanation for the lack of differences in recoil velocities. Increased amount of E-Cad also attests of stronger adhesive properties which act against tension created by the Myo-II. In any case, the proposed changes in mechanical forces in *aka* mutant cells remains such that it does not induce apical constriction and subsequent basal-cell extrusion or the induction of a fold in the tissue. The changes observed argue instead in favour of reinforcing adhesion to prevent cell extrusion.

Another argument in favour of a reinforcement of adhesive properties upon SJ alterations is the assembly of Vinc and Mys FAs laterally and basally. Although FA contacts, restricted to the basal site, are present at the location of attachment sites of flight muscles, rather late in pupal development [57], it is notable that FAs are being detected in *aka* clones as early as 15–16 h after puparium formation. Although Mys and Vinc are expressed in control epithelial cells, they do not necessarily assemble into FAs. Although we cannot exclude the possibility that the alteration of SJs may lead to transcriptional upregulation of Mys, we favour the hypothesis that reduced degradation of Mys by the ESCRT machinery contributes to FA assembly. We propose that such contacts contribute to the maintenance of epithelial cells within the epithelium layer, hence contributing to mutant cells’ preservation of epithelial mechanical integrity upon SJ disturbance.

Does a similar sensing mechanism exist in vertebrates upon alteration of TJs? A recent study revealed that serine proteinases are used to cleave the TJ complex form by proteins EpCAM and Claudin-7 upon TJ damages, releasing Claudin-7 and ensuring TJ rapid repair [60]. It remains to be determined whether such mechanisms described for small leaks apply to larger alterations of the TJ belt, as we report here in flies, and involve AJ and FA reinforcement of adhesive properties. Also, the observed changes at apical level might be mostly due to direct effects. Indeed, Tricellulin, a key tricellular TJ effector in vertebrates, has been shown to bind directly to the AJ component α-Catenin which then binds to F-actin and Vinculin. The binding of Tricellulin to α-Catenin is required to organize contractility and proper sealing at TCJ [61].

Due to their importance in ensuring epithelia homeostasis, deciphering between direct and indirect consequences of TJ alterations remains a key question to explore in the future.

## Acknowledgements

We thank A. Guichet, C. Klämbt, E. Knust, T. Lecuit, S. Luschnig, J. Mathieu, J. Treisman, K. Röper and A. Uv for reagents. We also thank J. R. Huynh and J. Mathieu (CIRB, Paris) for sharing the ShrubGFP CRISPR line prior to publication and the Bloomington Stock Center, the Vienna Drosophila RNAi Center and the National Institute of Genetics Fly Stock Center for providing fly stocks. We also thank S. Dutertre and X. Pinson from the Microscopy Rennes Imaging Center-BIOSIT (France). We are grateful to A. Dupont and A. Jankowska for excellent technical support for the electron microscopy. The JEM 2100 HT transmission electron microscope was available at the Laboratory of Microscopy, Department of Cell Biology and Imaging, Institute of Zoology and Biomedical Research, Jagiellonian University. The monoclonal antibodies against Cora and DE-Cad were obtained from the Developmental Studies Hybridoma Bank, generated under the auspices of the National Institute of Child Health and Human Development, and maintained by the University of Iowa Department of Biological Sciences.

## Author contributions

Conceptualization: T.E.d.B and R.L.B.; methodology: T.E.d.B, M.K.J., E.D and R.L.B.; investigation: T.E.d.B, M.K.J., E.D and R.L.B.; formal analysis: T.E.d.B and R.L.B.; visualization, T.E.d.B, M.K.J. and R.L.B.; writing – original draft: T.E.d.B and R.L.B.; writing – review and editing: T.E.d.B and R.L.B.; funding acquisition: T.E.d.B, M.K.J. and R.L.B.; supervision: R.L.B.

## KEY RESOURCES TABLE

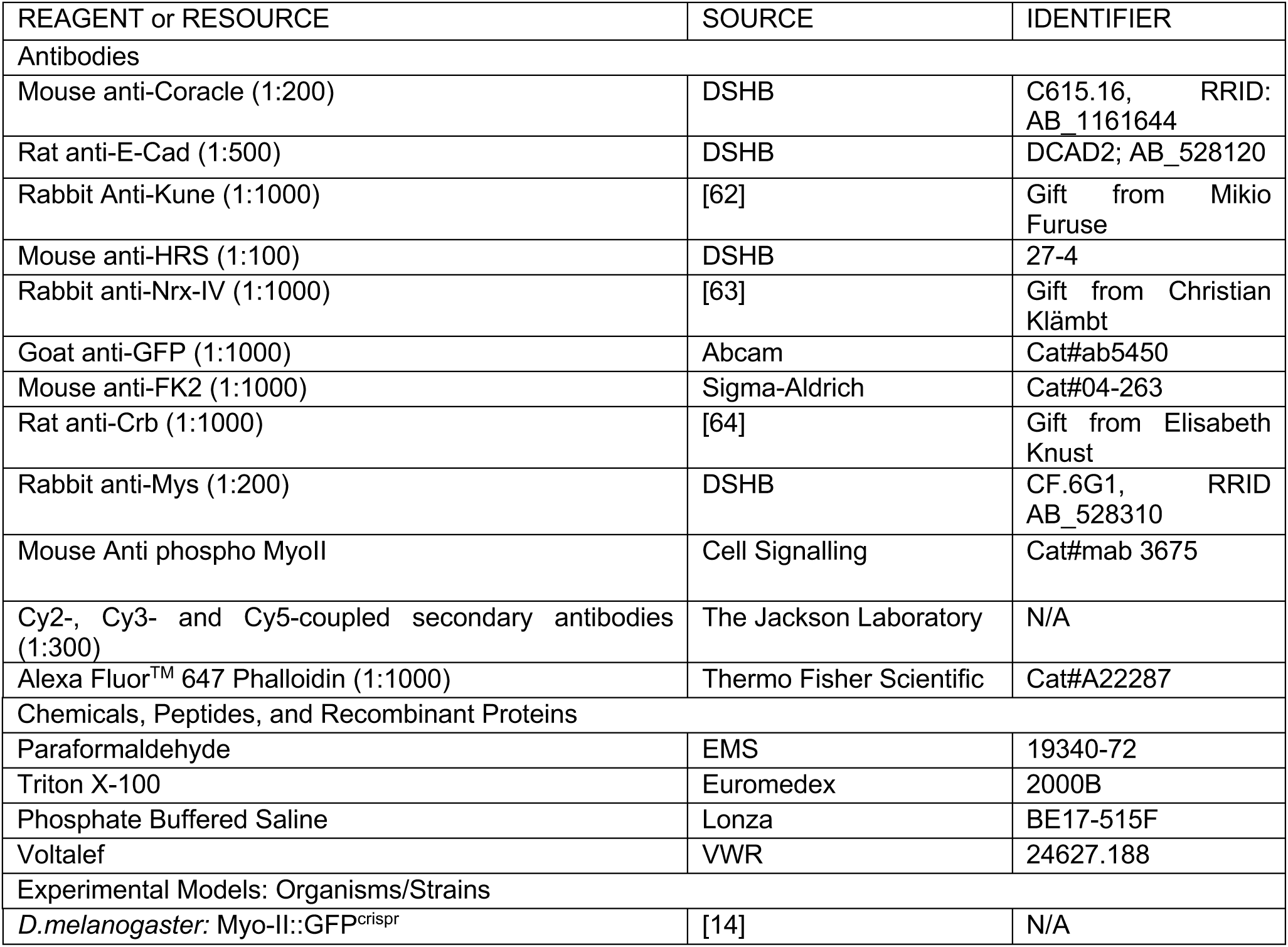
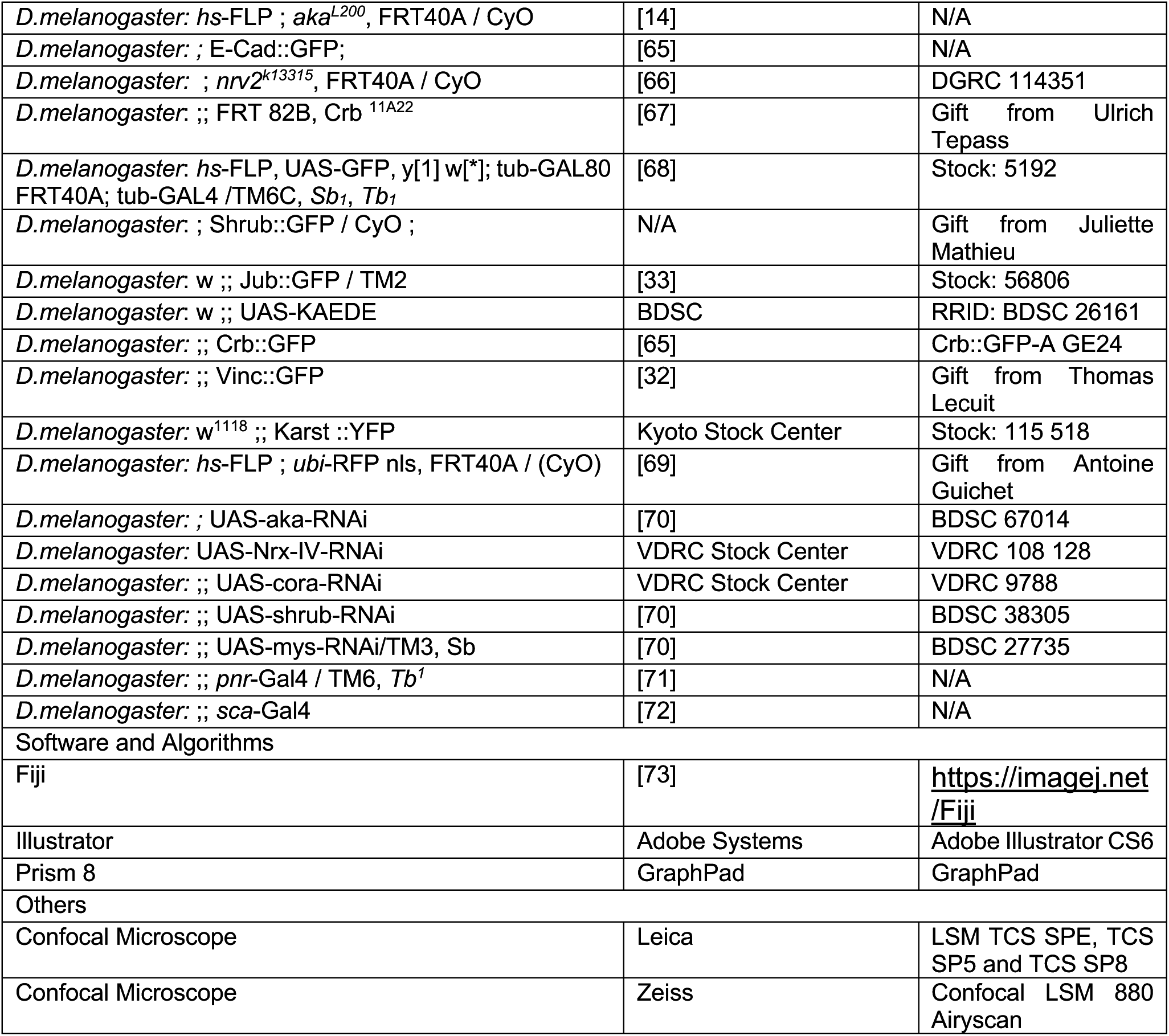

## Experimental model

### Drosophila genotypes

**Figure 1**

(A–A′) UAS-aka-RNAi ; *pnr*-Gal4 obtained by crossing UAS-aka-RNAi with *pnr*-Gal4 / TM6, *Tb1*

(B–B′) *hs*-FLP ; *aka^L200^*, FRT40A / *ubi*-RFP nls, FRT40A, E-Cad::GFP ; obtained by crossing *hs*-FLP ; *aka^L200^*, FRT40A / CyO with *hs*-FLP ; *ubi*-RFP nls, FRT40A, ECad::GFP / CyO

(D–D′) *hs*-FLP ; Myo-II::GFP; *ubi*-RFP nls, FRT40A / CyO obtained by crossing Myo-II::GFP; *ubi*-RFP nls, FRT40A / CyO ; with *hs*-FLP ; *aka^L200^*, FRT40A / CyO

(F–F′) *hs*-FLP ; *aka^L200^*, FRT40A / *ubi* -RFP nls, FRT40A obtained by crossing *hs*-FLP; *aka^L200^*, FRT40A / CyO with *hs*-FLP ; *ubi*-RFP nls, FRT40A / (CyO)

**Figure 2**

(A–A′ and C) *hs*-FLP ; Myo-II::GFP; *ubi*-RFP nls, FRT40A / CyO obtained by crossing Myo-II::GFP; *ubi*-RFP nls, FRT40A / CyO ; with *hs*-FLP ; *aka^L200^*, FRT40A /CyO

(F-G) *hs*-FLP ; *aka^L200^*, FRT40A / *ubi*-RFP nls, FRT40A, E-Cad::GFP ; obtained by crossing *hs*-FLP ; *aka^L200^*, FRT40A / CyO with *hs*-FLP ; *ubi*-RFP nls, FRT40A, ECad::GFP / CyO

Figure 3

(A-D’) Shrub::GFP ; UAS-cora-RNAi / UAS-KAEDE, *pnr*-Gal4 obtained by crossing UAS-cora-RNAi with ; Shrub::GFP ; UAS-KAEDE, *pnr*-Gal4 / SM5-TM6, *Tb*

Figure 4

(A) *sca*-Gal4 / UAS-shrub-RNAi obtained by crossing ;; *sca*-Gal4 with ;; UAS-shrub-RNAi

(B–C′′) Shrub::GFP ; UAS-cora-RNAi /*pnr*-Gal4 obtained by crossing UAS-cora-RNAi with ; Shrub::GFP ; *pnr*-Gal4 / SM5-TM6, *Tb*

(D–D′′) Shrub::GFP ; UAS-Nrx-IV-RNAi / *pnr*-Gal4 obtained by crossing UAS-Nrx-IV-RNAi with Shrub::GFP ; *pnr*-Gal4 / TM6, *Tb*

(E) *hs*-FLP ; *aka^L200^*, FRT40A / *ubi*-RFP nls, FRT40A obtained by crossing *hs*-FLP; *aka^L200^*, FRT40A / CyO with *hs*-FLP ; *ubi*-RFP nls, FRT40A / (CyO)

Figure 5

(A–A′′) hs-FLP ; *aka^L200^*, FRT40A / *ubi*-RFP nls, FRT40A ; Crb::GFP / + obtained by crossing *ubi*-RFP nls, FRT40A / CyO ; Crb::GFP / TM6, *Tb1* with *hs*-FLP ; *aka^L200^*, FRT40A / CyO

(B–B′′) *hs*-FLP ; *aka^L200^*, FRT40A / *ubi* -RFP nls, FRT40A obtained by crossing *hs*-FLP; *aka^L200^*, FRT40A / CyO with *hs*-FLP ; *ubi*-RFP nls, FRT40A / (CyO)

(D-D’) *hs*-FLP ; *aka^L200^*, FRT40A / *ubi*-RFP nls, FRT40A, E-Cad::GFP ; FRT82B, Crb^11a22^ / + obtained by crossing ; *aka^L200^*, FRT40A / CyO ; FRT82B, Crb^11a22^ / TM6, *Tb1* with *hs*-FLP ; *ubi*-RFP nls, FRT40A, ECad::GFP / CyO

Figure 6

(A–D) *hs*-FLP ; *akaL200*, FRT40A / *ubi* -RFP nls, FRT40A obtained by crossing *hs*- FLP ; *akaL200*, FRT40A / CyO with *hs*-FLP ; *ubi*-RFP nls, FRT40A / (CyO)

(E-E’) *hs*-FLP, UAS-GFP; *aka^L200^*, FRT40A / *tub*-GAL80, FRT40A; UAS-mys-RNAi- / *tub*-GAL4 obtained by crossing *aka^L200^*, FRT40A; UAS-mys-RNAi /SM5-TM6b, *Tb* with *hs*-FLP, UAS-GFP, *tub*-GAL80, FRT40A; *tub*-GAL4 /TM6C, *Sb*, *Tb*

Figure S1

(A–A′′) *hs*-FLP ; *nrv2^k13315^*, FRT40A / *ubi* -RFP nls, FRT40A, E-Cad::GFP ; obtained by crossing *hs*-FLP ; *nrv2^k13315^*, FRT40A / CyO with *hs*-FLP ; *ubi*-RFP nls, FRT40A, E-Cad::GFP / CyO

(C–C′′) *hs*-FLP / Myo-II::GFP ; *nrv2^k13315^*, FRT40A / *ubi*-RFP nls, FRT40A obtained by crossing Myo-II::GFP; *ubi*-RFP nls, FRT40A / CyO ; with *hs*-FLP ; *nrv2^k13315^*,FRT40A / CyO

(E-E’) *hs*-FLP ; *aka^L200^*, FRT40A / *ubi*-RFP nls, FRT40A, E-Cad::GFP ; obtained by crossing *hs*-FLP ; *aka^L200^*, FRT40A / CyO with *hs*-FLP ; *ubi*-RFP nls, FRT40A, ECad::GFP / CyO

Figure S2

(A–D′) hs-FLP ; *aka^L200^*, FRT40A / *ubi*-RFP nls, FRT40A ; Vinc::GFP / + obtained by crossing *ubi*-RFP nls, FRT40A / CyO ; Vinc::GFP / TM6, *Tb1* with *hs*-FLP ; *aka^L200^*, FRT40A / CyO

(E–E′) *hs*-FLP ; *aka^L200^*, FRT40A / *ubi*-RFP nls, FRT40A ; Karst::YFP / + obtained by crossing ; *ubi*-RFP nls, FRT40A / CyO ; Karst::YFP / TM6, *Tb1* with *hs*-FLP ; *aka^L200^*, FRT40A / CyO

(G-G’) *hs*-FLP ; *aka^L200^*, FRT40A / *ubi*-RFP nls, FRT40A ; Jub::GFP / + obtained by crossing ; *ubi*-RFP nls, FRT40A / CyO ; Jub::GFP / TM2 with *hs*-FLP ; *aka^L200^*, FRT40A / CyO

Figure S3

(A–A′) *hs*-FLP ; *aka^L200^*, FRT40A / *ubi* -RFP nls, FRT40A obtained by crossing *hs*-FLP ; *aka^L200^*, FRT40A / CyO with *hs*-FLP ; *ubi*-RFP nls, FRT40A / (CyO)

(C–C′) *hs*-FLP ; *nrv2^k13315^*, FRT40A / *ubi* -RFP nls, FRT40A ; obtained by crossing *hs*- FLP ; *nrv2^k13315^*, FRT40A / CyO with *hs*-FLP ; *ubi*-RFP nls, FRT40A / CyO

Figure S4

(A-B’’’’) *sca*-Gal4 / UAS-shrub-RNAi obtained by crossing ;; *sca*-Gal4 with ;; UAS- Shrub-RNAi

Figure S5

(A–A′′) *hs*-FLP ; *nrv2^k13315^*, FRT40A / *ubi*-RFP nls, FRT40A ; Crb::GFP / + obtained by crossing ; *ubi*-RFP nls, FRT40A / CyO ; Crb::GFP / TM6, *Tb1* with *hs*-FLP ; *nrv2^k13315^*, FRT40A / CyO

(B–B′′) *hs*-FLP ; *nrv2^k13315^*, FRT40A / *ubi* -RFP nls, FRT40A ; obtained by crossing *hs*- FLP ; *nrv2k13315*, FRT40A / CyO with *hs*-FLP ; *ubi*-RFP nls, FRT40A / CyO

Figure S6

(A-A’’’’) s*ca*-Gal4 / UAS-shrub-RNAi obtained by crossing ;; *sca*-Gal4 with ;; UAS- shrub-RNAi

## Method details

### Immunofluorescence

Pupae aged from 16h30 to 19h after puparium formation (APF) were dissected using Cannas microscissors (Biotek, France) in 1X Phosphate-Buffered Saline (1X PBS, pH 7.4) and fixed 15 min in 4% paraformaldehyde at room temperature [74]. Following fixation, dissected nota were permeabilized using 0.1% Triton X-100 in 1X PBS (PBT), incubated with primary antibodies diluted in PBT for 2 hours at room temperature (see the Key Resources Table for details of antibodies and dilutions used). After 3 washes of 5 minutes in PBT, nota were incubated with secondary antibodies in PBT for 1 hour, followed by 2 washes in PBT, and one wash in PBS, prior mounting in 0,5% N-propylgallate dissolved in 90% glycerol/PBS 1X final.

### Live-imaging and image analyses

Live imaging was performed on pupae aged for 16h30 APF at 25°C. Pupae were sticked on a glass slide with a double-sided tape, and the brown pupal case was removed over the head and dorsal thorax using microdissecting forceps. Pillars made of 4 and 5 glass coverslips were positioned at the anterior and posterior side of the pupae, respectively. A glass coverslip covered with a thin film of Voltalef 10S oil is then placed on top of the pillars such that a meniscus is formed between the dorsal thorax of the pupae and the glass coverslip [75]. Images were acquired with an LSM Leica SPE, SP5 or SP8 equipped with a 63X N.A. 1.4. objective and controlled by LAS AF software or by LSM Zeiss 880 AiryScan equipped with a 63X N.A.1.4. objective and controlled by ZEN software. Confocal sections (z) were taken every 0.5 µm or 1µm. For figures representation, images were processed with Gaussian Blur σ = 1.1. All images were processed and assembled using Fiji software [73] and Adobe Illustrator.

### Nanoablation

Laser ablation was performed on live pupae aged for 16h to 19h APF using a Leica SP5 confocal microscope equipped with a 63X N.A. 1.4 objective or a LSM Zeiss 880 AiryScan equipped with a 63X N.A. 1.4 objective. Ablation was carried out on epithelial cell membranes at AJ level with a two-photon laser-type Mai-Tai HP from Spectra Physics set to 800 nm and a laser power of 2.9W.

## Quantification and statistical analysis

### Fluorescence signal analysis

Sum slices were applied to different experiments. A circular ROI of 2µm*2µm was drawn to measure signal at vertices, a circular ROI of 3µm*3µm for the medial network and centered in the measured cells and a segmented line of 10 pixels width was used to measure signals at bicellular junctions. Using the same width or diameter, lines and circular ROI were drawn to extract background fluorescence signals and the background signal was subtracted to each quantification. After, data were standardized between 0 and 10 to allow visual representation with 10 corresponding to the highest signal in each experiment analyzed and 0 the lowest. Standardization was operated on data of cells belonging to the same *notum* in every experiment.

### Cell area quantification

Sum slices projection was applied then WT and *aka^L200^*cells were discriminated on their presence/absence of nls::RFP signal. We excluded cells at the border of the WT/ *aka^L200^* clonal area. A mask was applied based on the E-Cad::GFP or E-Cad signal and area in µm^2^ was extracted. Appropriate statistical tests were used to check for significative differences.

### Measurement of the length of the new adhesive interface at cytokinesis

The time t = 0 was set according to the frame just before the beginning of the contraction of the cell. Each frame was separated by 2 min. The maximal expected size of the junction was inferred shorly after the beginning of the contraction with the presumptive localization of the two future vertices, i.e. the place where the initial decrease of E-cad is detected [28, 30]. Then, at every time point, the length was measured at the new vertices formed and standardized to the initial maximal expected size.

### Statistical tests

All information concerning the statistical details are provided in the main text and in figure legends, including the number of samples analyzed for each experiment. Prism 8 software or R 4.2.1 were used to perform the analyses.

Scattered plots use the following standards: thick line indicate the means and errors bars represent the standard deviations. Boxplots with connected line use the following standards: dots represent mean and the total-colored areas show SD.

The Shapiro-Wilk normality test was used to confirm the normality of the data and the F-test to verify the equality of SD. The statistical difference of data sets was analyzed using the Student unpaired two-tailed t test, Multiple t tests, Fisher t test or the non-parametric Wilcoxon-Mann-Whitney test. Statistical significances were represented as follow: p value > 0.05 NS (not significant), p-value≤0.05*; p-value≤0.01** and p value ≤ 0.0001 ****.

