## Supplemental Figures for "ESCRT-III-dependent adhesive and mechanical changes are triggered by a mechanism sensing paracellular diffusion barrier alteration in *Drosophila* epithelial cells"

### Supplemental Figures, Esmangart de Bournonville et al

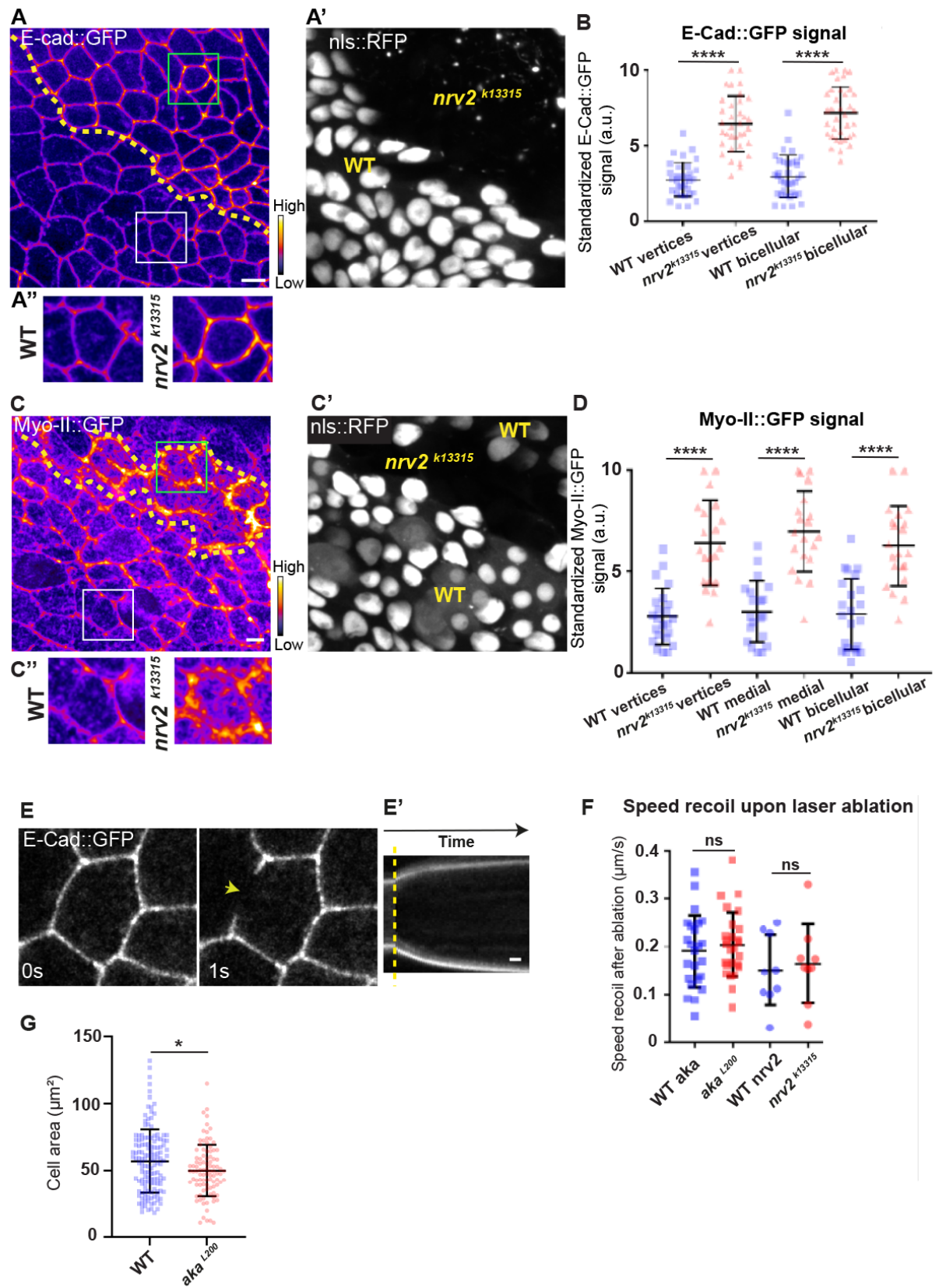

**Figure S1: Consequence of loss of Nervana2 and Anakonda on E-Cad and MyoII localization, and on cell-cell junction mechanical properties; related to main Figure 1**

(A–A') Localization of E-Cad::GFP (A, fire colour) or Myo-II::GFP (B, fire colour) in wild-type (A', C', nls::RFP-positive cells in grey) and *nrv2<sup>k13315</sup>* mutant cells (A', C' nls::RFP negative). Wild-type and *nrv2<sup>k13315</sup>* cells are separated by the dashed yellow lines in (A–C'). Higher magnification of yellow dashed square depicted in panels A and C for wild-type and *nrv2<sup>k13315</sup>* cells, respectively (A'' and C''). (B) Plot of the standardized E-Cad::GFP signal at tricellular and bicellular junctions in wild-type (blue squares) and *nrv2<sup>k13315</sup>* cells (red triangles) (n = 33 and 35 vertices and n = 35 and 36 bicellular junctions for wild-type and *nrv2<sup>k13315</sup>*, respectively; 2 pupae for each condition). (D) Plot of the standardized Myo-II::GFP signal at bicellular junctions, vertices as well as medial network in wild-type (blue squares) and *nrv2<sup>k13315</sup>* cells (red triangles) (n = 23 and 20 vertices and n = 20 cellular medial networks and n = 23 and 21 bicellular junctions for wild-type and *nrv2<sup>k13315</sup>*, respectively; n = 2 pupae for each condition). (E) Example of wild-type laser-based nanoablation in the AJ plane identified using E-Cad::GFP. Yellow arrowhead shows the nanoablation area. (E') Kymograph of the ablation area depicted in panel E, showing vertices' recoil upon ablation. Scale bar shows 5 s. (F) Plot of the mean recoil velocities upon nanoablation for wild-type (blue squares, n > 20 ablations, n > 5 pupae; circles, n = 9 ablations, n = 3 pupae) and *aka<sup>L200</sup>* (red squares, n > 20 ablations, n > 5 pupae) or *nrv2<sup>k13315</sup>* cells (red circles, n = 9 ablations, n = 3 pupae), respectively. (G) Quantification of the cell area (in $\mu\text{m}^2$) of WT (blue squares, n = 137 cells, n > 5 pupae) and *aka<sup>L200</sup>* cells (red circles, n = 96 cells, n > 5 pupae). Bars show mean ± SD, \*\*\*\*p < 0.0001, unpaired t test for panel B, D and F. \* p < 0.05, Welch's t test for panel G. A calibration bar shows LUT for grey value range. The scale bars represent 5 $\mu\text{m}$ for panels A and C. White squares represent close-up of WT and green squares of *nrv2<sup>k13315</sup>* situations for panels A and C.

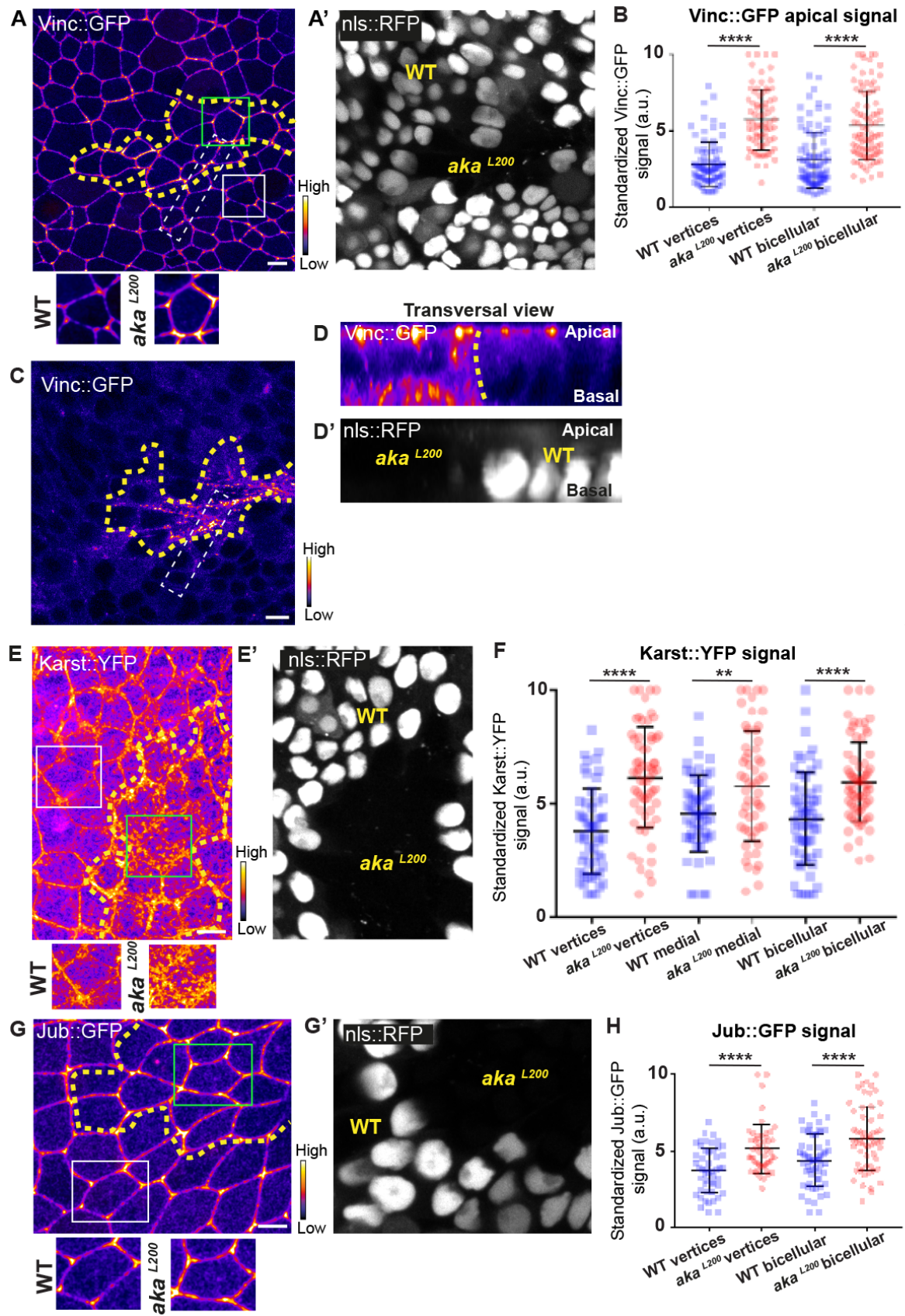

**Figure S2: Loss of Anakonda leads to enrichment of Vinculin, Karst and Ajuba at bi and tri cellular junctions; related to main Figure 2**

(A, C and D) Localization of Vinc::GFP in both wild-type and *aka<sup>L200</sup>* cells, at apical (A) and basal level (C) in a planar view or in a transversal view (D), separated by the dashed yellow line. (A' and D') Clones of wild-type and *aka<sup>L200</sup>* cells identified by nls::RFP marking. (B) Plot of the standardized Vinc::GFP signal at tricellular and bicellular junctions in wild type (blue squares) and *aka<sup>L200</sup>* cells (red circles). (n = 84 and 76 vertices and n = 92 bicellular junctions for wild-type and *aka<sup>L200</sup>* respectively, n = 4 pupae for each condition). (E) Localization of Karst::YFP in both wild-type and *aka<sup>L200</sup>* cells, separated by the dashed yellow line. (E') Clones of wild-type and *aka<sup>L200</sup>* cells identified by nls::RFP marking. (F) Plot of the standardized Karst::YFP signal at tricellular and bicellular junctions as well as medial network in wild type (blue squares) and *aka<sup>L200</sup>* cells (red circles). (n = 54 and 64 vertices and n = 55 and 56 cellular medial networks and n = 59 and 68 bicellular junctions for wild-type and *aka<sup>L200</sup>* respectively, n = 4 pupae for each condition). (G) Localization of Jub::GFP (fire color) in both wild-type and *aka<sup>L200</sup>* cells, separated by the dashed yellow line. (G') Clones of wild-type and *aka<sup>L200</sup>* cells identified by nls::RFP marking. (H) Plot of the standardized Jub::GFP signal at tricellular and bicellular junctions in wild type (blue squares) and *aka<sup>L200</sup>* cells (red circles). (n = 50 vertices and n = 60 bicellular junctions for wild-type and *aka<sup>L200</sup>* respectively, n = 2 pupae for each condition). Bars show Mean ± SD, \*\*\*\* p < 0.0001, unpaired t test and Mann-Whitney test. A calibration bar shows LUT for grey value range. White dotted rectangle on A shows area depicted in D-D'. The scale bars represent 5μm. White squares represent close-up of WT and green squares of *aka<sup>L200</sup>* situations for panels A, E and G.

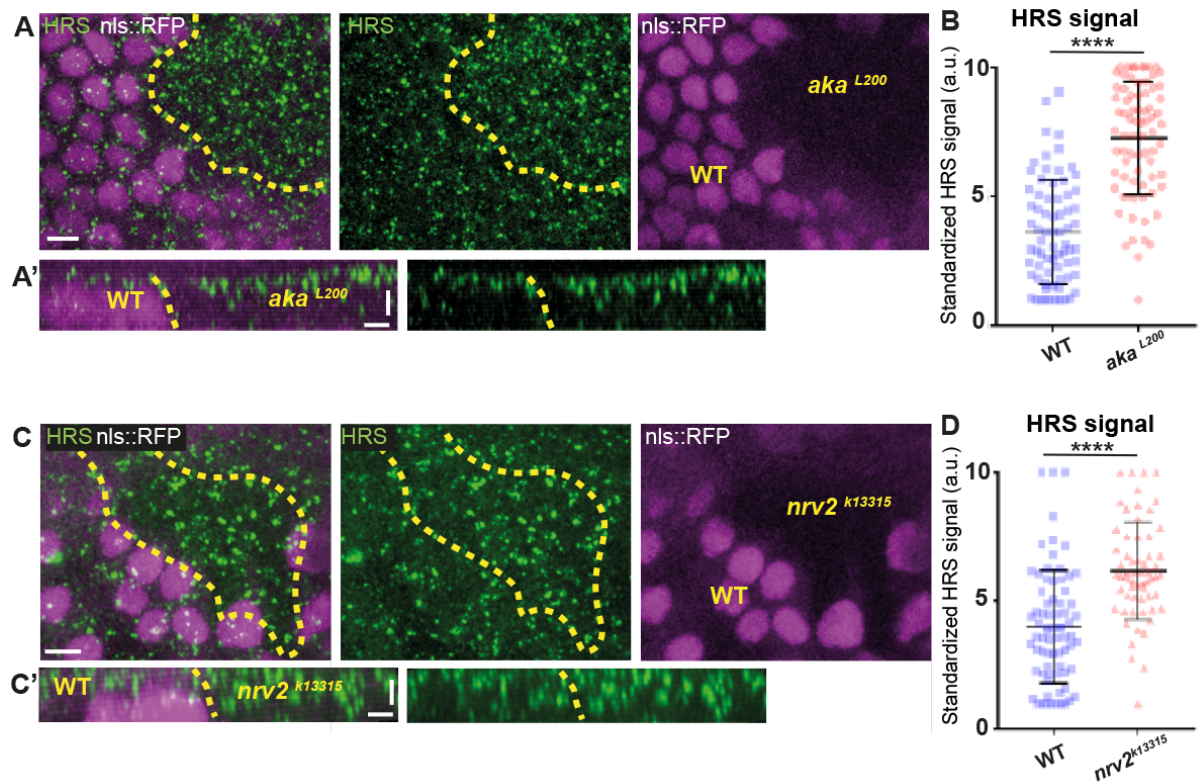

**Figure S3: Loss of Anakonda or Nervana 2 triggers increased number of HRS-positives vesicles; related to main Figure 3**

(A) Localization of HRS (green) in wild-type and *aka*<sup>L200</sup> cells, separated by the dashed yellow line. Clones of wild-type and *aka*<sup>L200</sup> cells identified by nls::RFP marking (magenta). (A') Transversal view of (A). (B) Plot of the standardized HRS signal wild type (blue squares) and *aka*<sup>L200</sup> cells (red circles). (n = 72 and n = 75 for wild-type and *aka*<sup>L200</sup> cells respectively, n > 5 pupae for each condition). (C) Localization of HRS (green) in wild-type and *nrv2*<sup>k13315</sup> cells, separated by the dashed yellow line. Clones of wild-type and *nrv2*<sup>k13315</sup> cells identified by nls::RFP marking (magenta). (C') Transversal view of (C). Plot of the standardized HRS signal wild type (blue squares) and *nrv2*<sup>k13315</sup> cells (red triangle). (n = 74 and n = 62 for wild-type and *nrv2*<sup>k13315</sup> cells respectively, n = 5 pupae for each condition). Bars show Mean ± SD, \*\*\*\* p < 0.0001, unpaired t test and Mann-Whitney test.

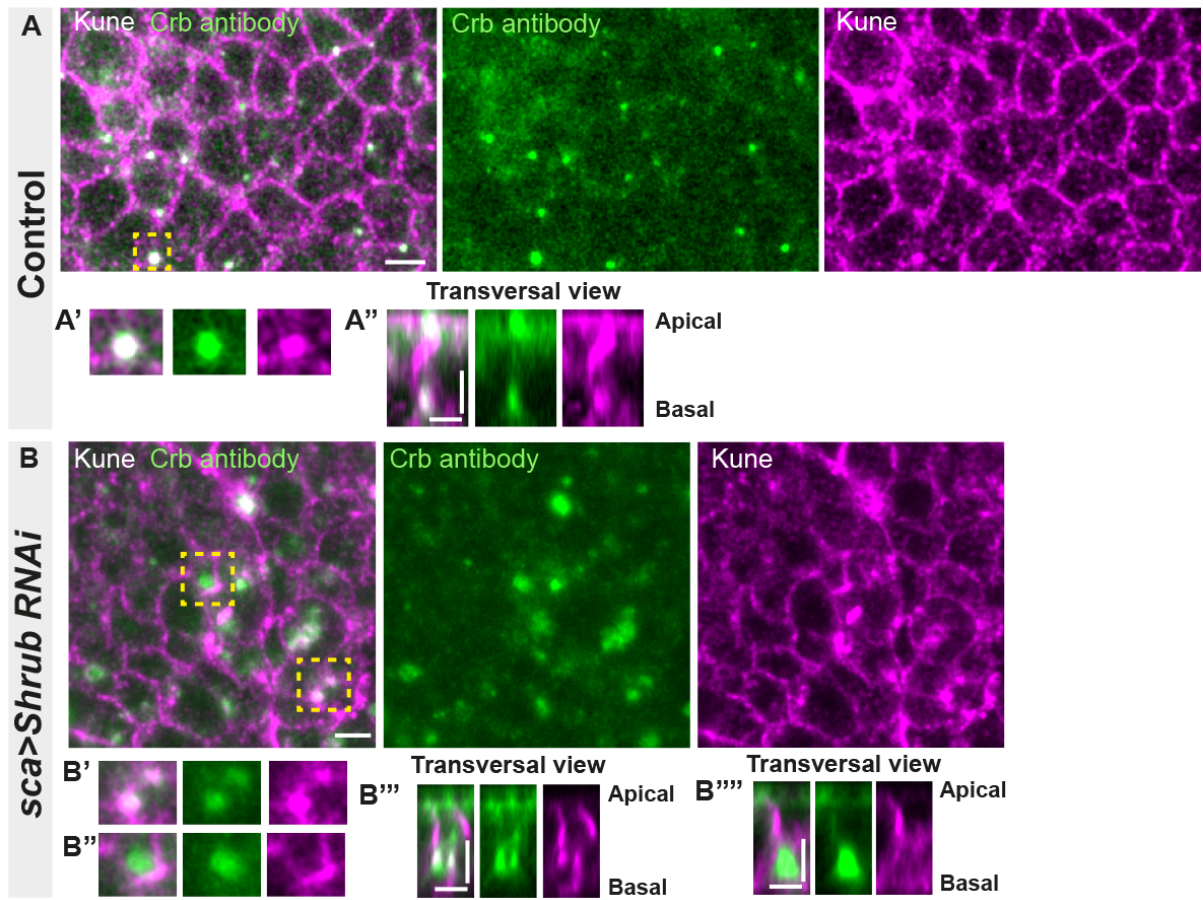

**Figure S4: Loss of function of ESCRT III protein Shrub in *notum* cells leads to Crumbs and septate junction protein Kune abnormal localization; related to main Figure 5**

(A-A'') Localization of Crb (anti Crb, green) in a wild-type area of cells expressing UAS::*shrub*-RNAi under *sca*-Gal4 control and marked by SJ protein Kune (anti Kune, magenta). Yellow dashed square shows (A' and A'') magnification of cells with colocalization between Crb and Kune in vesicles at basal cell level in a planar view (A') and transversal view (A''). (B-B''') Localization of Crb (anti Crb, green) in cells expressing UAS::*shrub*-RNAi under *sca*-Gal4 control and marked by SJ protein Kune (anti Kune, magenta, Kune disrupted signal). Yellow dashed square shows (B'-B''') magnification of cells with partial (B') or no colocalization (B'') between Crb and Kune at basal cell level in a planar view (B'-B'') and transversal view (B'''-B'''). The scale bar represents 5 $\mu\text{m}$ (A and B) and 3 $\mu\text{m}$ in (A'', B'' and B''').

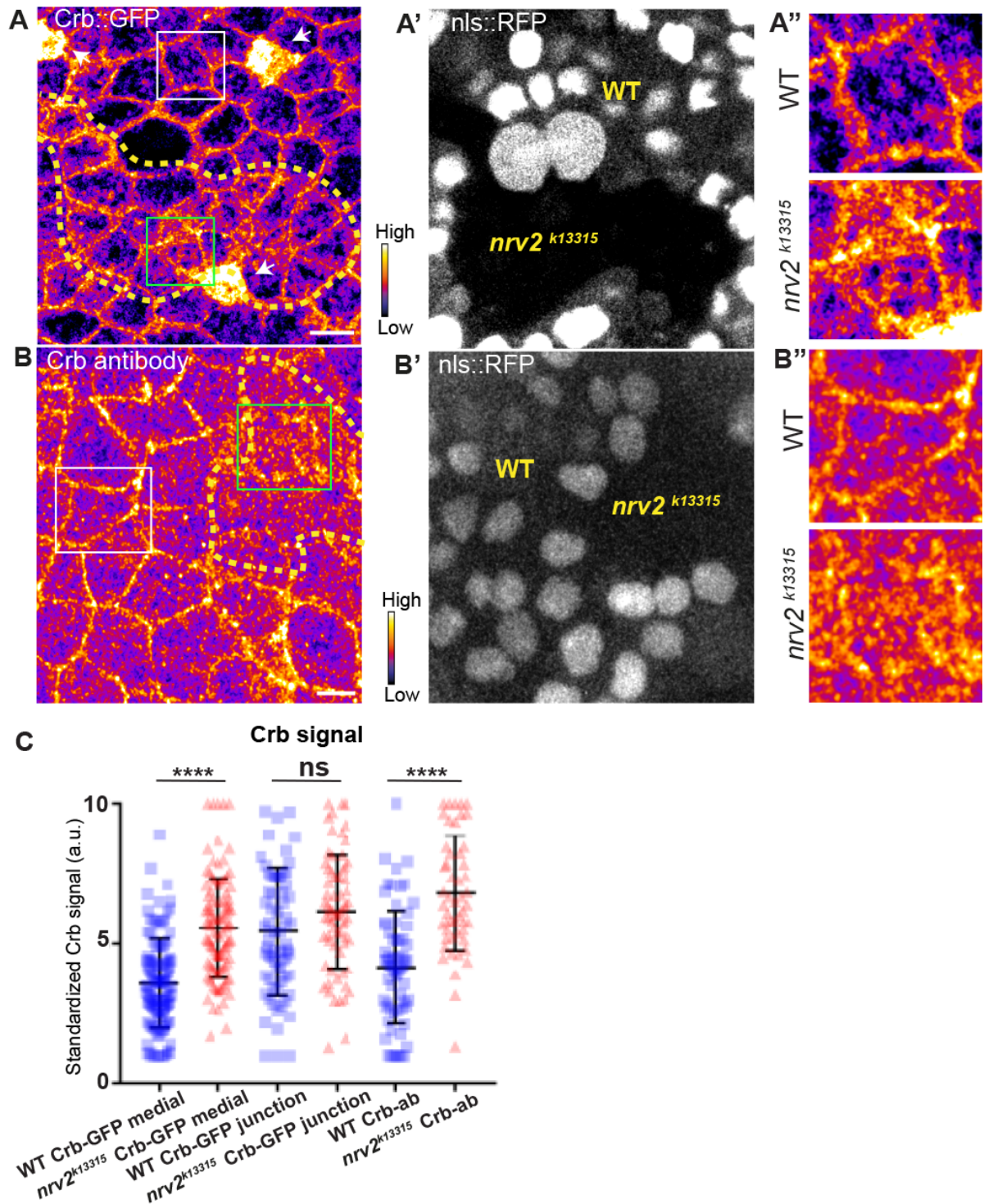

**Figure S5: Loss of Nervana 2 leads to higher level of Crumbs at adherens junction level; related to Figure 5**

(A-A') Localization of Crb::GFP in both wild-type and *nrv2*<sup>k13315</sup> cells, separated by the dashed yellow line. (A') Clones of wild-type and *nrv2*<sup>k13315</sup> cells identified by nls::RFP marking. White squares show (A'') magnification of cells depicted in (A). (B-B'') Localization of anti-Crb in both wild-type and *nrv2*<sup>k13315</sup> cells, separated by the dashed yellow line. (B') Clones of wild-type and *nrv2*<sup>k13315</sup> cells identified by nls::RFP marking. White squares show (B'') magnification of cells depicted in (B). (C) Plot of the standardized Crb::GFP signal at the medial and junctional part of the cell or Crb-ab only at the medial part, in wild type (blue squares) and *nrv2*<sup>k13315</sup> cells (red triangles). (n = 75 cellular medial networks, n = 70 junctions and n = 68 and 52 cellular medial networks with Crb-ab for wild-type and *nrv2*<sup>k13315</sup> respectively, n = 3 pupae for Crb::GFP and n > 5 pupae for Crb-ab). Bars show Mean ± SD, \*\*\*\* p < 0.0001, Mann-Whitney test. A calibration bar shows LUT for grey value range. The scale bars represent 5µm. White squares represent close-up of WT and green squares of *nrv2*<sup>k13315</sup> situations for panels A and B.

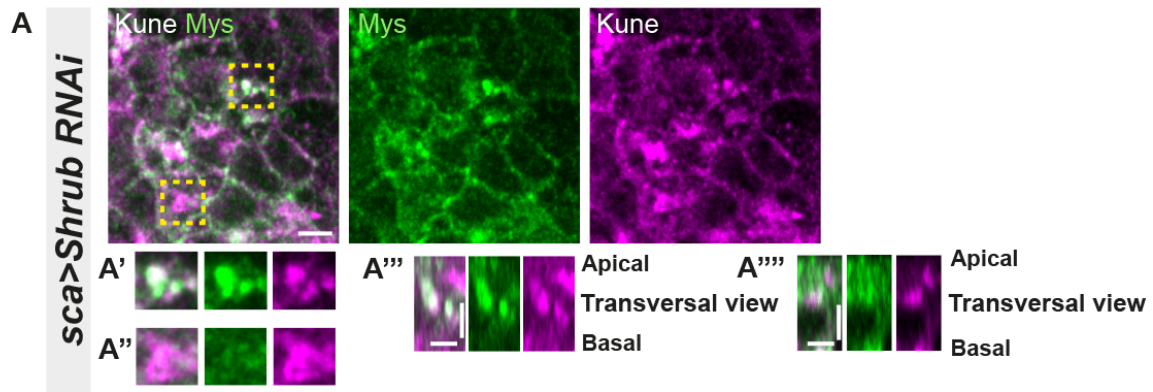

**Figure S6: Loss of function of Shrub in *notum* cells leads to Myospheroid and Kune-Kune abnormal localization; related to Figure 6**

(A-A'') Localization of Mys (anti-Mys, green) and Kune (anti-Kune, magenta) in cells expressing UAS::shrub-RNAi under sca-Gal4 control. Yellow dashed square shows (A'-A'') magnification of cells with (A') or without colocalization (A'') between Mys and Kune at basal cell level in a planar view (A'-A'') and transversal view (A'''-A''). The scale bar represents 5μm (A) and 3μm in (A''' and A'').
